# Neural determinants of the increase in muscle strength and force steadiness of the untrained limb following a 4-week unilateral training

**DOI:** 10.1101/2025.03.25.645159

**Authors:** E. Lecce, P. Amoruso, A. Del Vecchio, A. Casolo, F. Felici, D. Farina, I. Bazzucchi

## Abstract

Enhanced untrained muscle strength and force steadiness following unilateral resistance training (i.e., *cross-education*) is typically attributed to neural responses. However, the mechanisms of these adaptations for spinal motoneurons remain unexplored. Therefore, we examined maximal-voluntary-force (MVF), steady-force variability (CovF), and longitudinally tracked motor unit adaptations in 10 individuals completing a 4-week unilateral strength intervention compared to 9 controls. High-density surface electromyography was recorded from the biceps brachii during steady (10%MVF) and trapezoidal (35%MVF) contractions. The relative proportion of common synaptic input (CSI) to motoneurons and its variability (CSI-V) were estimated using coherence and spectral analysis. Indirect estimates of persistent inward currents using firing rate hysteresis (ΔF) and motor unit recruitment thresholds (MURT) were assessed during the ramp forces (35%MVF). MVF increased in both the trained (+14%, *p*<0.001) and untrained limbs (+6%, *p*=0.004), and CovF decreased in both limbs (*p*<0.001). Greater CSI was observed on both sides (p<0.01), concomitant with reduced CSI-V (p<0.01). ΔF increased exclusively in the trained limbs (+1.61±0.71 pps; *p*<0.001), and both sides exhibited lower MURT (p<0.001). In trained limbs, MVF gains were strongly associated with changes in CSI, MURT, and ΔF (R²>0.70, *p*<0.01), while the contralateral muscle MVF increase was associated exclusively with CSI and MURT (R²>0.65, *p*<0.01). In both limbs, lower CovF was strongly associated with reduced CSI-V (R²>0.70, *p*<0.01). Our findings suggest that enhanced untrained muscle force and steadiness are mediated by increased relative strength of shared synaptic input with respect to independent noise and decreased variability of this shared input, with gains in trained muscle MVF being associated with ΔF.

**Key points:**

- Unilateral resistance training improves strength and force steadiness in the contralateral untrained limb, suggesting neural adaptations without directly overloading the muscle.
- Despite established force-related modifications, the specific neural mechanisms mediating these functional responses remain largely unknown.
- 4-week unilateral training intervention enhanced muscle strength and force steadiness in the untrained limbs of 10 individuals, alongside a greater proportion of shared synaptic input, reduced variance in common input, and lower motor unit recruitment thresholds.
- We demonstrate that the neural mechanisms underlying improved strength and force control in muscles without mechanical overloading are associated with a higher relative shared input among motoneurons and reduced variance in these common input components.

Graphical abstract legend
unilateral training intervention reduced the variance in common synaptic input (CSI-V), which was associated with decreased variability in force steadiness (CovF) in both the trained and contralateral untrained limbs. On the exercised side, the increase in maximal voluntary force (MVF) was accompanied by a higher proportion of common synaptic input (CSI), a lower motor unit recruitment threshold (RT), enhanced persistent inward current (PIC) amplitude, and increased neural drive. In contrast, the contralateral untrained limb exhibited higher shared synaptic input and a lower RT but with unaltered PIC amplitude and neural drive (⇋). Overall, these adaptations resulted in a 14% increase in MVF in the exercised limb and a 6% increase in the untrained limb.

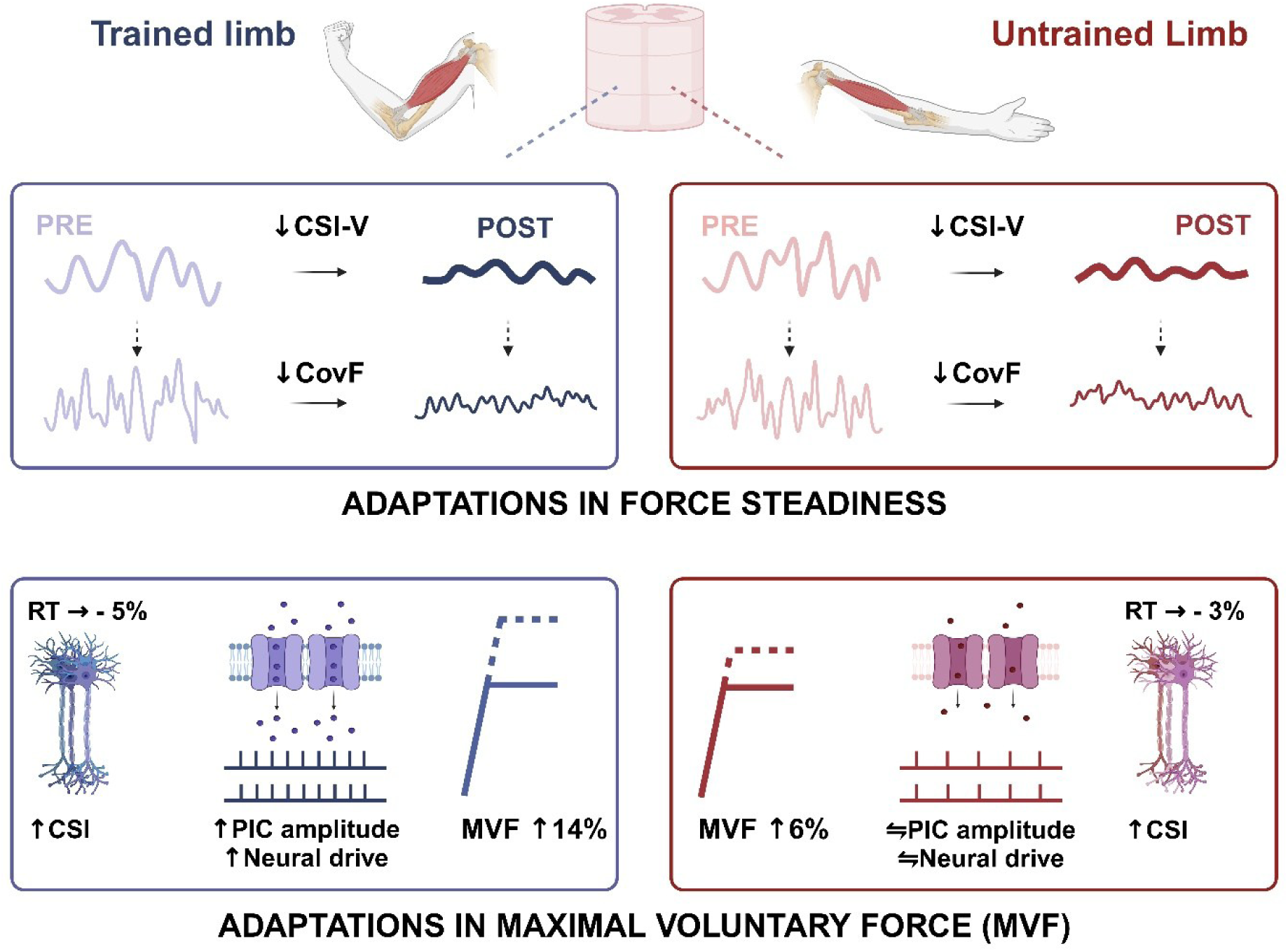

## Introduction

It is well documented that resistance training elicits significant strength and skill enhancements through multiple neural and morphological modifications (Del Vecchio et al., 2024; Folland & Williams, 2007; Green & Gabriel, 2018a; Škarabot et al., 2021; Vila-Chã & Falla, 2016). Gains within the first 4 weeks are believed to be predominantly mediated by neural mechanisms (Del Vecchio, Casolo, et al., 2019) with minimal contribution from contractile muscle adaptations, as evidenced by unaltered muscle architecture and twitch properties in this training interval (Blazevich et al., 2007; Carroll et al., 2009; Škarabot et al., 2021; Van Cutsem et al., 1998). The high relevance of neural adaptations is also reflected by the interesting observation that *unilateral* resistance training typically results in increased muscle strength and force accuracy also for the untrained contralateral homologous muscle, a phenomenon termed cross-education (*cross-transfer* or *interlimb transfer*) (Green & Gabriel, 2018a). On average, the untrained limb exhibits a 7% increase in force output, approximately half the gain observed on the trained side (Hendy & Lamon, 2017). However, the strength gains largely depend on the intervention type and duration, with eccentric contractions inducing up to 20% of force increase by interlimb transfer (Frazer et al., 2018; Green & Gabriel, 2018a; Hendy & Lamon, 2017; Latella et al., 2012; Manca et al., 2018; Sato et al., 2021). Because of the lack of muscle loading, contralateral gains in function with unilateral training must be due exclusively to neural adaptations.

*Interlimb transfer* has been extensively explored and associated with multiple spinal and supraspinal mechanisms (Carson, 2020; Hendy & Lamon, 2017; Manca et al., 2018; Pearcey et al., 2022; Ruddy & Carson, 2013). Notably, we have recently demonstrated that enhanced maximal voluntary force (MVF) and steady force accuracy in untrained limbs were accompanied by motor unit adaptations, including a lower recruitment threshold, increased dynamic range of discharge rate from the onset to its peak, and a reduced variability in the interspike intervals (Lecce, Conti, et al., 2025). In the current study, we focused on common synaptic input to motoneurons and intrinsic motoneuron properties, and we studied their associations with improvements in force steadiness and maximal force following contralateral muscle training.

The relative strength of common vs independent input to motoneurons can be quantified by coherence analysis (A. M. Castronovo, Negro, & Farina, 2015; Farina et al., 2013; Hug, Del Vecchio, et al., 2021; Negro et al., 2016) and determines the level of synchronisation in motoneuron discharges as well as the variability in discharges. This relative strength, however, does not directly impact force variability because the independent input has a negligible effect on the neural drive to the muscle and, therefore, on force variability. Conversely, the absolute variance (or power) of the common synaptic input has a direct effect on force variability (A. Castronovo et al., 2018; J. L. Dideriksen et al., 2012; Feeney et al., 2018). In addition to the input they receive, the output of motoneurons (neural drive to muscle) also depends on the intrinsic properties of motoneurons, which are modulated by persistent inward currents (PICs) (Heckman et al., 2005; Orssatto et al., 2023).

In this study, we examined the neural responses to a 4-week unilateral resistance training intervention by longitudinally tracking motor units in the biceps brachii of both the trained and untrained limbs. Building on our previous investigation (Lecce, Conti, et al., 2025), we employed a single tracking procedure (pre-post) and contractions at 10% and 35% MVF to enable a proper exploration of the shared synaptic input and estimates of PICs. We expected an increase in maximal force and a decrease in force variability on both sides following unilateral training. Given the known relations between maximal force and force steadiness with common synaptic input to motoneurons and motoneuron intrinsic properties, we hypothesised that (*a*) the relative strength of the common input would increase and its absolute variance decrease during steady contraction phases following training, (*b*) intrinsic excitability of motoneurons would increase by neuromodulatory pathways and (*c*) these adaptations would be associated with relative changes in the maximal strength and force steadiness.

## Methods

### Participants and ethical approval

The study was approved by the local ethics committee of the University of “Foro Italico,” Rome (approval - CAR157/2023), and adhered to the standards outlined in the Declaration of Helsinki. All participants provided written informed consent, which detailed the experimental procedures and potential risks, emphasising their right to withdraw from the study at any time without jeopardy. A total of 20 recreationally active participants (10 males and 10 females) were initially recruited. One participant withdrew, leaving 19 participants who completed the protocol. Following a familiarisation session, participants were randomly assigned to either the intervention group (INT, n = 10; females = 5) or the control group (CNT, n = 9; females = 4) using a block-randomisation approach to ensure equal group sizes (Kang et al., 2008). Each participant was assigned a unique alphanumeric code to protect their privacy and confidentiality. Exclusion criteria included metabolic diseases, upper limb musculoskeletal disorders, acute infections, uncontrolled hypertension, use of medications that affect muscle protein metabolism, vascular tone, or neural activity, and use of contraceptives following previous longitudinal setup in resistance training (Burrows & Peters, 2007; Elliott-Sale et al., 2020; Lecce, Romagnoli, et al., 2024; Reif et al., 2021). Inclusion criteria required participants to be between 18 and 35 years old and in good health.

### Overview of the study

Participants attended three laboratory visits. The first visit was dedicated to familiarisation, and the subsequent two were for neuromuscular testing conducted at baseline (Pre) and after resistance training (Post). Volunteers in the intervention group completed 12 training sessions (three per week) over four weeks, with a washout period of five to seven days before neuromuscular testing to minimise muscle soreness and residual training effects (Cheung et al., 2003; Mizumura & Taguchi, 2024).

During the first visit, participants were informed about the experimental procedures and prepared for neuromuscular testing by familiarising themselves with maximal voluntary isometric contractions (MVICs), isometric trapezoidal contractions, and steady contractions of the biceps brachii, which was selected due to its primary role in generating elbow flexion force (Dartnall et al., 2008; B. Yu et al., 2022). Additionally, the biceps brachii has been extensively investigated in cross-education research (Boyes et al., 2017; Green & Gabriel, 2018b; Howatson et al., 2011; Martinez et al., 2021; Sato et al., 2021) and is known to exhibit significant cross-transfer adaptations (Farthing & Chilibeck, 2003; Lecce, Conti, et al., 2025; Manca et al., 2018).

Participants’ dominant limb was identified using the Edinburgh Handedness Inventory Questionnaire (Oldfield, 1971), and the non-dominant limb was designated for the training intervention to maximise the observable cross-education effects. No measurements were taken during the first visit. Data collection took place in the second visit (Pre), during which assessments of MVF, trapezoidal and steady isometric contractions, and high-density surface electromyography (HDsEMG) recordings were obtained from the biceps brachii of both limbs (Fig 1A). The same data collection was repeated following the intervention or control period (Post).

**Figure 1.**
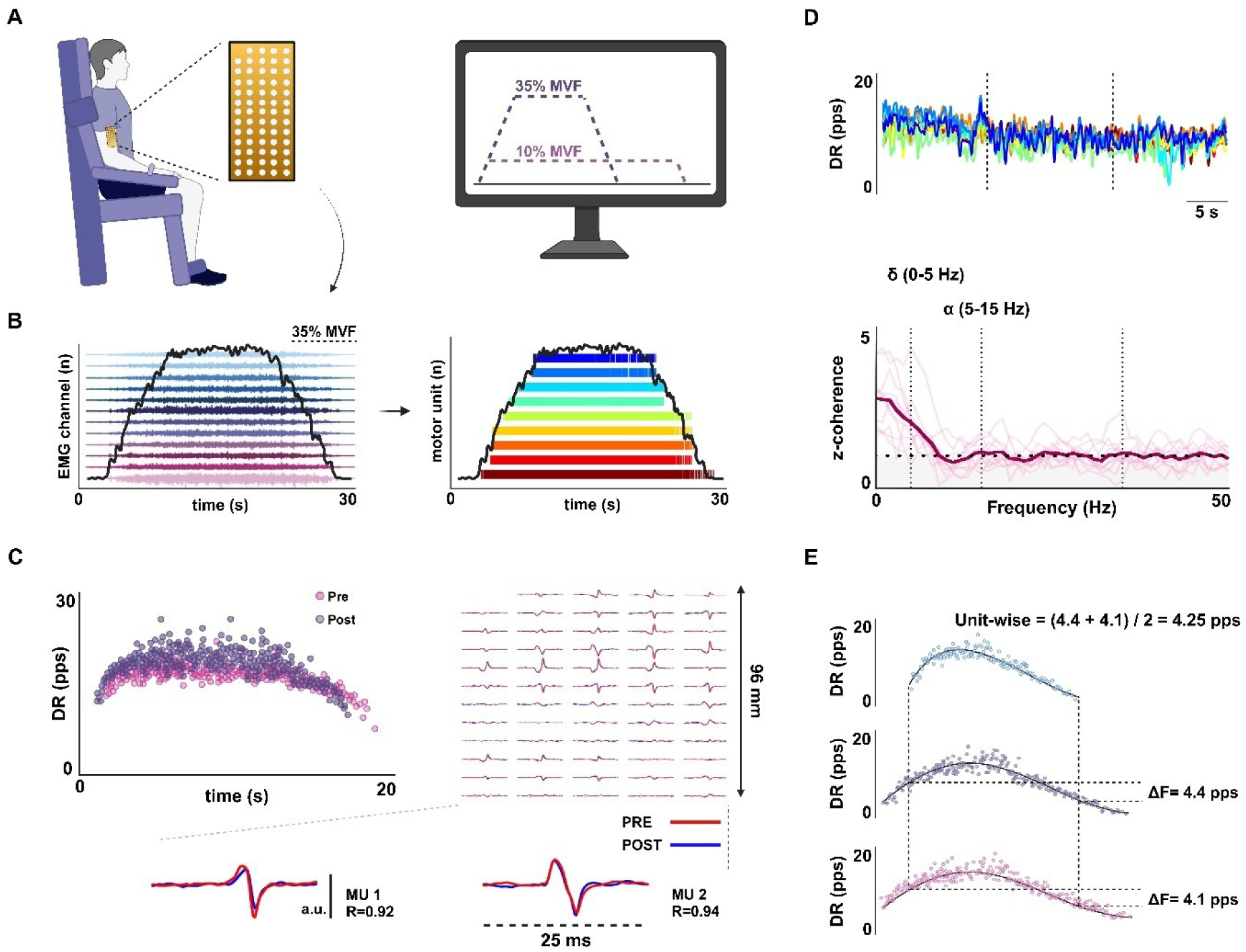
Overview of the study and EMG - decomposition analysis. A, Experimental setup for neuromuscular assessment. A bi-dimensional electrode grid (13 × 5) was placed over the biceps brachii to record myoelectrical activity during 10% and 35% MVF contractions. Participants sat on a dynamometric chair with the wrist secured in a cuff attached to a load cell for force recording. B, Monopolar HDsEMG recordings were synchronised with force output (black line) and decomposed into motor unit spike trains (colour-coded raster plot), where each row represents the discharge times of a single motor unit. C, Instantaneous discharge rate of the same motor unit tracked pre- and post-intervention based on waveform (2 dimensional) cross-correlation (cut-off: R > 0.8). Data from two motor units (MU1 and MU2) are shown. D, Smoothed discharge rate over time, illustrating window selection for coherence analysis. The central portion was chosen to ensure a stable discharge rate. Coherence was computed using Welch’s periodogram with 1 s non-overlapping windows, comparing the same motor units before and after the intervention. Z-scores were obtained for δ (0 - 5 Hz) and α (5 - 15 Hz) bandwidths, with bias defined as the mean coherence between 100 - 250 Hz. The significance threshold was set at the 95% confidence limit. E, The paired motor unit method estimated firing rate hysteresis (ΔF) in valid unit pairs. If multiple pairs were considered valid, unit-wise ΔF was calculated as the sum of ΔF values divided by the number of pairs.

For female participants, testing was scheduled during either the ovulatory or mid-luteal phase to minimise fluctuations in neuromuscular performance and ensure consistent hormonal influence across participants (Lecce, Conti, et al., 2024; Tenan et al., 2013; Weidauer et al., 2020). These phases were chosen due to the similarity of low-threshold motor unit firing rates during ovulation and the mid-luteal phase in order to standardise testing under conditions of heightened hormonal influence while avoiding the variability associated with the early follicular phase (Piasecki et al., 2023). The menstrual cycle was tracked using the validated Menstrual Practices Questionnaire (MPQ), which collects detailed information on menses onset, average cycle length, and regularity (Hennegan et al., 2020). Day one of the cycle was defined as the first day of menstrual bleeding. Based on self-reported data, forward counting was used to estimate the ovulatory and mid-luteal phases, ensuring phase alignment between test-retest sessions. Participants recorded the onset and duration of their last two cycles to confirm cycle regularity and accurately predict subsequent phases. This non-invasive, standardised approach ensured that testing sessions were aligned with specific menstrual phases. Post-intervention testing was conducted during the same menstrual phase as baseline to maintain consistency (Piasecki et al., 2024).

### Experimental procedures

The volunteers were instructed to refrain from strenuous exercise and caffeine consumption for 24-48 hours before testing (Bazzucchi et al., 2011). Neuromuscular testing began with a standardised warm-up, consisting of three sets of 10 isometric biceps brachii contractions at a low intensity (30 - 40% of the perceived MVF), with 30 seconds of rest between sets. Participants were encouraged to focus on isolating biceps brachii activation during force generation. Following the warm-up, participants performed three MVICs with 180 seconds of rest between trials. Each trial required maximal effort for 5 seconds, with strong verbal encouragement to ensure optimal performance (Lecce, Romagnoli, et al., 2024; Romdhani et al., 2024). The MVF was determined as the peak value recorded across the three MVICs and was subsequently used to set the relative target force (%MVF) for submaximal contractions. Five minutes after the final maximal trial, participants completed a steady isometric contraction at 10% of MVF for 60 seconds, followed by a trapezoidal contraction at 35% of MVF. This trapezoidal contraction consisted of a ramp-up phase at a force increase rate of 5% MVF·s^-1^, a 10-second plateau phase, and a ramp-down phase at a force decrease rate matching the ramp-up phase. Submaximal relative forces (10% and 35% of MVF) were set before and after the intervention, with values adjusted according to the MVF measured in each testing visit. This type of assessment is commonly employed in neuromuscular testing procedures assessing adaptations in motor unit discharge characteristics to resistance training (Del Vecchio, Casolo, et al., 2019; Orssatto et al., 2023), as using the same absolute force values before and after training could result in measurements taken at a lower percentage of the maximal force capacity post-training, potentially underestimating contraction intensity (Orssatto et al., 2023).

### Intervention and control protocols

The intervention group performed a 4-week unilateral resistance training protocol at the isokinetic dynamometer (Kin-com, Chattanooga, Tennessee), with three sessions per week using the *non-dominant limb*. This selection is based on evidence indicating that novel or less frequently performed motor tasks can elicit a greater cross-education effect, potentially owing to reduced habitual use and a lower baseline level of neuromuscular adaptation (Farthing et al., 2007). Although some studies report similar cross-education outcomes for dominant and non-dominant limb training (Song et al., 2024), we selected the non-dominant limb because it is less involved in daily activities, reducing potential interference with resistance training adaptations. Each session began with a standardised warm-up, including three sets of 10 dynamic contractions at 30% MVF, with 60 seconds of rest between sets. This was followed by two sets of four eccentric contractions at 50% MVF, with 120 seconds of rest between sets. Subsequently, participants performed four sets of six elbow-flexor eccentric contractions at a speed of 30°/s (from 140° to 40° of flexion) at 80% MVF, with 180 seconds of rest between sets. Control group participants did not perform any intervention and were instructed to maintain their habits throughout the experimental period. This approach ensured that no behavioural modifications occurred, allowing any observed changes in the experimental groups to be attributed solely to the intervention.

### Force signal recording

Elbow-flexion force was measured using an isokinetic dynamometer (load cell - Kin-Com, Chattanooga, Tennessee). Participants were seated in the dynamometric chair and stabilised with chest and waist straps to minimise extraneous movement. The upper arm was positioned parallel to the trunk, while the forearm was oriented midway between supination and pronation, maintaining a 90° elbow flexion angle. The centre of rotation of the lever arm was aligned with the distal lateral humerus epicondyle, and the wrist was secured in a cuff attached to the load cell to ensure force isolation. The analogue force signal was amplified and sampled at 2048 Hz using an external analogue-to-digital (A/D) converter (EMG - 400, OT - Bioelettronica, Turin, Italy) to ensure synchronisation with the electromyogram. For steady and trapezoidal contractions requiring visual guidance, participants were provided with a steady or trapezoidal force pattern (10% and 35% MVF, respectively) displayed throughout the contraction. A ±2% MVF error margin was indicated along the trajectory to minimise deviations and facilitate maintenance of the desired force pattern (Fig 1A).

### High-density surface electromyographic recordings

Electromyographic recordings were conducted one limb at a time, randomising among participants, using a 64-electrode adhesive grid (13 rows × 5 columns; gold-coated; electrode diameter: 1 mm; inter-electrode distance [IED]: 8 mm; OT-Bioelettronica, Turin, Italy). Following skin preparation, which included shaving, light abrasion, and cleansing with 70% ethanol, the biceps brachii long head perimeter was identified through palpation and marked with a surgical pen. The grid orientation was determined based on preliminary recordings from a 16-electrode array (IED: 5 mm, OT-Bioelettronica, Turin, Italy), allowing identification of the innervation zone (IZ) to estimate fibre direction. The IZ was located by identifying the inversion point in the action potential propagation direction, both proximally (towards the biceps brachii proximal tendon) and distally (towards the distal tendons), along the electrode column. The grid was then positioned directly over the IZ on the muscle belly using a disposable biadhesive with layer holes adapted to the HDsEMG grids (SpesMedica, Battipaglia, Italy). These holes were filled with conductive paste (SpesMedica, Battipaglia, Italy) to ensure optimal skin-electrode contact. To ensure consistent electrode positioning across all assessment time points, anatomical landmarks and skin marks were traced onto individual acetate sheets during the first assessment session (Orssatto et al., 2023). Reference electrodes were placed on the ulna (styloid process) and the acromion skin surface. HDsEMG signals were recorded in monopolar mode and digitised using a 16-bit multichannel amplifier (EMG-Quattrocento, OT-Bioelettronica, Turin, Italy). Signals were amplified (×150), sampled at 2048 Hz, and band-pass filtered (10 - 500 Hz) before being stored for offline analysis.

### Force and HDsEMG processing and analysis

The force signal was converted to newtons (N), and gravity compensation was used to remove the offset. The signal was low pass filtered with a fourth-order, zero-lag Butterworth filter with a cut-off frequency of 15 Hz. Only contractions without any counter-movement action or pre-tension were analysed (Del Vecchio et al., 2018). Before decomposing into individual motor unit action potentials, the monopolar EMG recordings were band-pass filtered at 20 - 500 Hz using a second-order, zero-lag Butterworth filter. The raw HDsEMG signals were decomposed using a blind source separation method (Fig 1B), namely convolutive kernel compensation (CKC) (Holobar et al., 2014; Holobar & Zazula, 2007). CKC generates filters for individual motor units, which estimate the motor unit spike trains when applied to the HDsEMG signal (Holobar et al., 2014). This decomposition procedure enables the identification of motor unit discharge times across a broad range of forces (Holobar & Farina, 2014). All the offline analysis was performed using DEMUSE software within MATLAB (MathWorks Inc., Natick, United States).

An experienced investigator manually analysed all identified motor units, following procedures extensively described in previous studies (Del Vecchio, Holobar, et al., 2020; Hug, Avrillon, et al., 2021; Valli et al., 2024). Manual editing involved inspecting and editing motor unit spike trains after automatic decomposition, ensuring accuracy by discarding motor units with a PNR below the reference threshold (30 dB), an interspike interval < 25 ms or > 250 ms (instantaneous spike frequency > 40 Hz or < 4 Hz, respectively), and/or an interspike interval variability (ISIv) > 30% (Del Vecchio, Holobar, et al., 2020; Hug, Avrillon, et al., 2021; Pearcey et al., 2024; Škarabot et al., 2023; Valli et al., 2024). Motor unit duplicates, defined as units sharing at least 30% of the same discharge activity, were identified and removed based on a firing match tolerance of 0.5 ms, with the motor unit having the lower PNR discarded (Škarabot et al., 2023). The manual editing approach adopted for motor unit spike trains has been shown to be highly reliable across operators (Hug, Avrillon, et al., 2021).

Motor units were longitudinally tracked across the intervention (pre-post) to ensure reliable comparisons (Fig 1C). This tracking method, based on the cross-correlation of two-dimensional action potential waveforms (Martinez-Valdes et al., 2017), has been widely used in interventional studies to directly assess the same set of motor units (Del Vecchio, Casolo, et al., 2019; Lecce, Conti, et al., 2025; Orssatto et al., 2023). Only motor units with a high cross-correlation coefficient (arbitrary R > 0.8) were included in the analysis. Tracking procedures were performed between trials of the same contraction intensity (e.g., 35% MVF: Pre - Post) to ensure data reliability.

Average motor unit discharge rate (MUDR) was calculated as the number of discharges from the series of discharge times divided by the steady phase time window, representing the number of discharges per second for each motor unit throughout the relative activity period (Del Vecchio, Negro, et al., 2019). For trapezoidal contractions, MU-ISIv and the coefficient of variation of force (CovF), defined as the percent ratio between the standard deviation and the mean force ([SD/mean]x100), were computed over the central 8 seconds of the plateau phase, excluding the first and last 1-second periods to minimise fluctuations associated with the ramp-up and ramp-down transitions. MU-ISIv and CovF were also computed for the central 20- second segment of steady contractions. Motor unit recruitment thresholds (MURT), which is the force value corresponding to the first motor unit spike, were computed from the series of discharge times identified by the decomposition exclusively in trapezoidal contractions (during the ramp-up phase). The MUDR, MU-ISIv, and MURT were averaged per participant to obtain the mean discharge rate (DR), ISIv, and RT for each individual. All the above procedures were performed following recent motor unit analysis guidelines (Del Vecchio, Holobar, et al., 2020; Gallina et al., 2022; Martinez-Valdes et al., 2023; Valli et al., 2024) using custom-written MATLAB scripts.

### Coherence analysis

Coherence analysis was conducted between cumulative spike trains (CSTs) of motor units, calculated as the sum of discharge times from two motor units randomly selected from those identified within a muscle (Maillet et al., 2022) in order to estimate the relative strength of common synaptic input (Lecce, Del Vecchio, et al., 2025). Coherence quantifies the correlation between two signals at specific frequencies, ranging from 0 (no correlation) to 1 (perfect correlation). Specifically, CSTs were cross-correlated by incrementally grouping motor units (e.g., eight identified motor units were divided into two groups, each containing up to four units). For each iteration, all unique combinations of motor units were tested, with a maximum of 100 random permutations (Maillet et al., 2022). Coherence was computed using Welch’s periodogram with non-overlapping 1-s Hanning windows. This approach enabled coherence estimation across various contraction intensities and window durations using controlled steady-force patterns of differing lengths (Cabral et al., 2024; Del Vecchio, Holobar, et al., 2020; Goodlich, Del Vecchio, Horan, et al., 2023). Pre-post comparisons were performed on the same contraction intensity and equal window length to ensure robust and reliable comparisons.

Coherence within the δ (0 - 5 Hz) band reflects the presence of the common drive (De Luca & Erim, 1994; Hug et al., 2023), while coherence within the α (alpha, 5 - 15 Hz) band indicates contributions from muscle afferences and spinal circuitries (Farina et al., 2017; Williams & Baker, 2009). Reliable coherence estimation in the β (15 - 35 Hz) band requires a larger motor unit sample due to the non-linear relationship between synaptic input to motoneurons and their output signal. This non-linearity is more pronounced at higher frequencies due to the undersampling of synaptic input (Farina & Negro, 2015; Negro & Farina, 2012). Consequently, coherence within the β band is not considered in this study.

This analysis was conducted exclusively within the central 20-second window of 1-minute steady contractions at 10% MVF and the central 8 seconds of the plateau phase in trapezoidal contractions at 35% MVF, excluding the 1-second transitions from ramp to plateau and the entire ramp-up and ramp-down portions (Cabral et al., 2024; Lecce, Del Vecchio, et al., 2025; McManus et al., 2019). These phases were discarded because of the large modulation variability of the discharge rate during the increasing/decreasing force portion compared to a steady force pattern, in which a more stable discharge rate is expected (S. N. Baker et al., 1997; Kakuda et al., 1999; Kilner et al., 1999). Indeed, a weak correlation of the motor units firing is typically observed during asynchronous recruitment, whereas a higher degree of correlation is obtained when motor units fire together due to shared synaptic input from the central nervous system (Del Vecchio, Sylos-Labini, et al., 2020). Furthermore, correlations between neural spike trains are influenced by the frequency at which action potentials are evoked, with higher spike frequencies implicating a greater degree of correlation (De La Rocha et al., 2007). Consequently, coherence was estimated during steady-force patterns, characterised by constant discharge rates, to relatively match the instantaneous discharge rates of motor units (Del Vecchio, Sylos-Labini, et al., 2020).

Previous studies have demonstrated that the correlation between cumulative motoneuron discharge trains increases monotonically with the total number of discharges included in the calculation of each CST (A. Castronovo et al., 2018; A. M. Castronovo, Negro, Conforto, et al., 2015; Nuccio et al., 2024). The rate of this increase depends on the proportion of common input relative to independent input, with a steeper increase observed when common input is more prevalent (A. M. Castronovo, Negro, Conforto, et al., 2015). Consequently, when a similar number of discharges (or the same motor units in a longitudinal setting) are used in the estimation, differences in coherence reflect differences in the strength of common input (Nuccio et al., 2024). To ensure robustness, the same set and number of motor units were used for coherence calculations throughout longitudinal tracking, allowing differences in coherence to reflect variations in the strength of common input.

Coherence values (C) were transformed into standard z-scores by first converting them to Fisher’s values [FZ= atanh 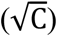] and then normalising to the variance of estimation [ZC=FZ / 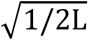], where L represents the number of segments used to calculate coherence without overlapping (e.g., 20 windows of 1 s, L = 20). This normalisation enhances the robustness and reliability of comparisons (A. Castronovo et al., 2018). Estimated bias was calculated as the mean coherence values within the 100 - 250 Hz range, where no significant coherence is expected, and subtracted from coherence profiles (A. Castronovo et al., 2018; Nuccio et al., 2024). Significant threshold coherence was defined as z-scores exceeding 0.83 at 10% MVF and 1.37 at 35% MVF, representing the 95% confidence limit (Maillet et al., 2022). The mean and maximal coherence values were calculated within the δ and α frequency bandwidth (Fig. 1D).

The proportion of common synaptic input relative to the total synaptic input received by motoneurons was estimated as the rate of increase in low-frequency coherence with increasing motor unit numbers (Hug, Del Vecchio, et al., 2021; Negro et al., 2016). This proportion reflects the shared synaptic input among motoneurons, referred to as the neural signal selectively amplified by the motoneuron pool, acting as a spatial filter and transmitting this input as neural drive to muscles (Farina et al., 2014; Negro & Farina, 2011). Low-frequency oscillations of the neural drive were estimated by low-pass filtering the composite spike train with a 400-ms Hanning window and detrending the resulting signal. The variability of these oscillations, termed ‘common noise,’ was quantified as the standard deviation of the detrended signal (i.e., the square root of the signal power). This metric reliably reflects the fluctuations in the common synaptic input to motoneurons (CSI-V), as previously indicated (A. Castronovo et al., 2018).

### Estimates of the persistent inward currents

PICs were estimated using a custom MATLAB script to calculate motor unit onset-offset hysteresis (ΔF) from pairs of motor units identified by correlating their discharge rates (Gorassini et al., 1998). Correlation was performed within consecutive 500-ms windows using the rate-rate correlation, an index of common modulation between two concurrently active motor units. The average discharge rate for each 500-ms bin was plotted for both the reference and test motor units throughout the trapezoidal contraction (Powers & Heckman, 2015). Only correlations with r ≥ 0.7 were considered, as lower values do not meet the assumption of shared synaptic input (Orssatto et al., 2021). Since PICs in the reference unit require up to 1.5 seconds to fully activate before the test motor unit is recruited, pairs with recruitment time differences of less than 1.5 seconds were excluded (Binder et al., 2020). Additionally, a test motor unit must be derecruited at least 1.5 seconds before the reference unit to ensure accurate ΔF calculations and avoid overestimation (Goodlich, Del Vecchio, Horan, et al., 2023). Control units exhibiting discharge rate saturation after the test unit recruitment (i.e., > 0.5 peaks per second) were discarded (Orssatto et al., 2023).

The discharge events for each motor unit were converted into instantaneous discharge frequencies and fitted with a fifth-order polynomial function (Orssatto et al., 2021). The earliest recruited motor unit was designated as the control, while the other was treated as the test unit (Gorassini et al., 1998). ΔF was calculated as the difference in discharge rate of the control motor unit at the times of recruitment and derecruitment of the test motor unit (Fig. 1E). When more than two motor units were identified for potential pairing, the sum of ΔF values was divided by the number of pairs [e.g., for three pairs= (ΔF_1_+ΔF_2_+ΔF_3_)/3] to obtain the *unit-wise* (Goodlich, Del Vecchio, Horan, et al., 2023).

Although simulation studies have suggested that steady phases of contraction could influence the estimation of PICs (Revill & Fuglevand, 2011), recent *in vivo* research has successfully employed trapezoidal contractions to obtain accurate estimates of PICs. These studies, adopting both iEMG and HDsEMG across a broad range of contraction intensities (10-50% MVF), have confirmed the sensitivity of this approach using both triangular and trapezoidal contractions (Goodlich, Del Vecchio, Horan, et al., 2023; Jenz et al., 2023; Martino et al., 2024; Mesquita et al., 2023, 2024; Orssatto et al., 2021, 2023; Wilson et al., 2015), by also confirming high test-retest reliability (Lapole et al., 2024). Therefore, unit-ΔF and participant-ΔF (i.e., averaged unit-ΔF for each individual) were computed using 35% MVF trapezoidal contractions.

By using longitudinally tracked data (matched motor units), only pairs of motor units identified at both time points were included. This approach ensures that the discharge rates of the same units are compared between timelines. Although these procedures significantly reduced the number of unit-ΔF compared to analysing the complete set of identified motor units, we characterised at least one ΔF value for each participant across limbs and timelines.

### Statistical analysis

The Shapiro-Wilk test was conducted to assess the normality of the data distribution, confirming that all data were normally distributed. The average number of identified motor units was evaluated using one-way ANOVA. Differences in MVF, CovF, mean and maximal z-coherence across all frequency bandwidths, as well as CSI-V, were analysed using paired-sample t-tests. This approach was chosen to account for related measurements within participants, considering the mean trend, thus enhancing statistical precision compared to one-way ANOVA or independent-sample t-tests, which assume independent observations (Cohen, 1988). Differences between the delta (%) MVF between limbs were also evaluated using paired-sample t-tests. Reliability and *agreement* in measurements were evaluated for decomposition-derived variables (MUDR, MU-ISIv, MURT, and ΔF) using two-way mixed-effects intraclass correlation coefficients (ICC_3,1_) using the *SimplyAgree* R-package, jamovi module. These analyses were employed to determine whether the decomposition process provides reliable absolute values despite the presence of an intervention (Ten Hove et al., 2024).

Differences in MUDR, MU-ISIv, MURT, and unit-ΔF over time were analysed using mixed-effects linear regression, preserving intra- and inter-participant variability by incorporating the entire sample of extracted motor units from the 19 participants (Wilkinson et al., 2023; Z. Yu et al., 2022). Fixed effects included group, limb, time, and their interactions, with a random intercept for each participant [e.g., MUDR ∼ group × limb × time + (1 | Participant ID)], following guidelines for the analysis of motor unit properties using this approach (Goodlich, Del Vecchio, Horan, et al., 2023; Héroux, 2021; Maillet et al., 2022; Orssatto et al., 2023; Wilkinson et al., 2023). Bonferroni corrections were applied for significant interactions, and estimated marginal means with 95% confidence intervals were computed for pre- and post-intervention comparisons. Paired-sample t-tests were also used to compare the effects of the intervention for DR, ISIv, RT, and participant-ΔF (Héroux, 2021).

The coefficient of determination (R²) was calculated to assess the association between averaged ΔF and DR, evaluating the strength of the relationship between the PIC-related gain control system and neural drive (Lee & Heckman, 1999, 2000; Orssatto et al., 2021). The strength of the association was interpreted as follows: 0 - 0.1, *very weak*; 0.1 - 0.3, *weak*; 0.3 - 0.5, *moderate*; 0.5 - 0.7, *strong*; 0.7 - 1.0, *very strong* (Ingene & Weisberg, 1981). In order to assess the influence of neural variables on muscle mechanical behaviour, associations between the changes in MVF with δ z-coherence - RT - ΔF - DR and the changes in CovF with CSI-V - ISIv, from baseline to post-intervention, were assessed. The robustness of this averaged approach was ensured through consistent tracking procedures (Del Vecchio, Negro, et al., 2019; Lecce, Conti, et al., 2025; Martinez-Valdes et al., 2017). To infer whether the neuromechanical adaptations imply a transfer to the contralateral untrained side, the association between the change in δ z-coherence, CSI-V, MVF, and CovF between limbs were also computed.

Cohen’s d was calculated as the effect size for significant results, reported with 95% confidence intervals (Goodlich, Del Vecchio, Horan, et al., 2023). Statistical analyses were performed using jamovi 2.3.28 (The jamovi project, Sydney, Australia) and SPSS 25.0 (IBM Corp., Armonk, NY, United States). A *p*-value of < 0.05 was considered statistically significant. The full statistical report, which includes INT and CNT results, is available as supplementary material.

## Results

### Anthropometrics, MVF, and CovF

Between-group comparisons of anthropometric characteristics and MVF at baseline are presented in Table 1.

**Table 1:** Baseline comparisons.

| Measurement | INT | CNT | *p* |
| --- | --- | --- | --- |
| Height (cm) | 168±9 | 172±7 | 0.190 |
| Weight (kg) | 62±12 | 70±19 | 0.266 |
| BMI (kg m<sup>-2</sup>) | 21.8±2.5 | 23±4.7 | 0.460 |
| Age (years) | 24.4±1.6 | 24.5±1.7 | 0.891 |
| MVF D (N) | 281±98 | 303±97 | 0.626 |
| MVF ND (N) | 261±89 | 272±93 | 0.789 |

CNT, control group; D, dominant limb; INT, intervention group; ND, non-dominant limb. Statistical comparisons were performed with independent sample t-tests. Data are presented as the mean±SD.

In the INT group, MVF significantly increased in both the trained (ΔMVF = +28 N [17, 40], ∼14%, *p* < 0.001, *d* = 1.79 [0.75, 2.80]) and untrained (ΔMVF = +17 N [7, 26], ∼6%, *p* = 0.004, *d* = 1.23 [0.38, 2.05]) limbs. Delta comparisons revealed a significantly higher degree of strength increase for the trained muscles compared to the contralateral untrained side (ΔMVF = 8% [4, 12], *p* = 0.002, *d* = 1.38 [0.48, 2.24]). In contrast, the CNT group showed no significant changes in MVF for either limb (*p* > 0.05). In trained limbs, CovF decreased at both 10% MVF (ΔCovF = -0.70% [0.38, 1.03], *p* < 0.001, *d* = 1.56 [0.60, 2.48]) and 35% MVF (ΔCovF = - 0.60% [0.46, 0.74], *p* < 0.001, *d* = 3.11 [1.55, 4.63]). Similarly, CovF was significantly lower at both 10% MVF (ΔCovF = -0.56% [0.35, 0.76], *p* < 0.001, *d* = 1.94 [0.85, 3.01]) and 35% MVF (ΔCovF = -0.51% [0.39, 0.63], *p* < 0.001, *d* = 3.12 [1.56, 4.65]) in the untrained limbs. The control group presented no changes in CovF for both limbs (*p* > 0.05).

### HDsEMG decomposition and tracking accuracy

Considering both groups, testing conditions and contractions, a total of 1266 motor units were identified following the decomposition process. 701 units were extracted from the INT group, with 354 at the baseline (per participant: TL, 17.8 ± 3.6 - UT, 17.6 ± 4.3) and 347 post-intervention (per participant: TL, 17.5 ± 3.2 - UT, 17.2 ± 3.8), while 565 units were identified in the CNT group with 292 at the baseline (per participant: D, 16.3 ± 1.1 - ND, 16.1 ± 1.2) and 273 post-intervention (per participant: D, 15.4± 2.2 - ND, 14.9 ± 1.9). Approximately 30% of the identified motor units were tracked across timelines and groups. Specifically, 200 units were tracked in the INT group (trained limb, *n* = 100; untrained limb, *n* = 100), and 175 units were tracked in the CNT group (non-dominant limb, *n* = 86; dominant limb, *n* = 89). On average, 20.0 ± 3.1 motor units were tracked per participant in the INT group (trained limb, 10.0 ± 2.3; untrained limb, 10.0 ± 1.8) and 19.4 ± 2.2 in the CNT group (non-dominant limb, 9.6 ± 1.1; dominant limb, 9.9 ± 1.3) across contractions. No significant differences were observed in the average number of identified motor units between groups and participants (*p* > 0.05). The number of identified and tracked motor units classified by limb, timeline, and contraction intensities is summarised in Table 2.

**Table 2:**
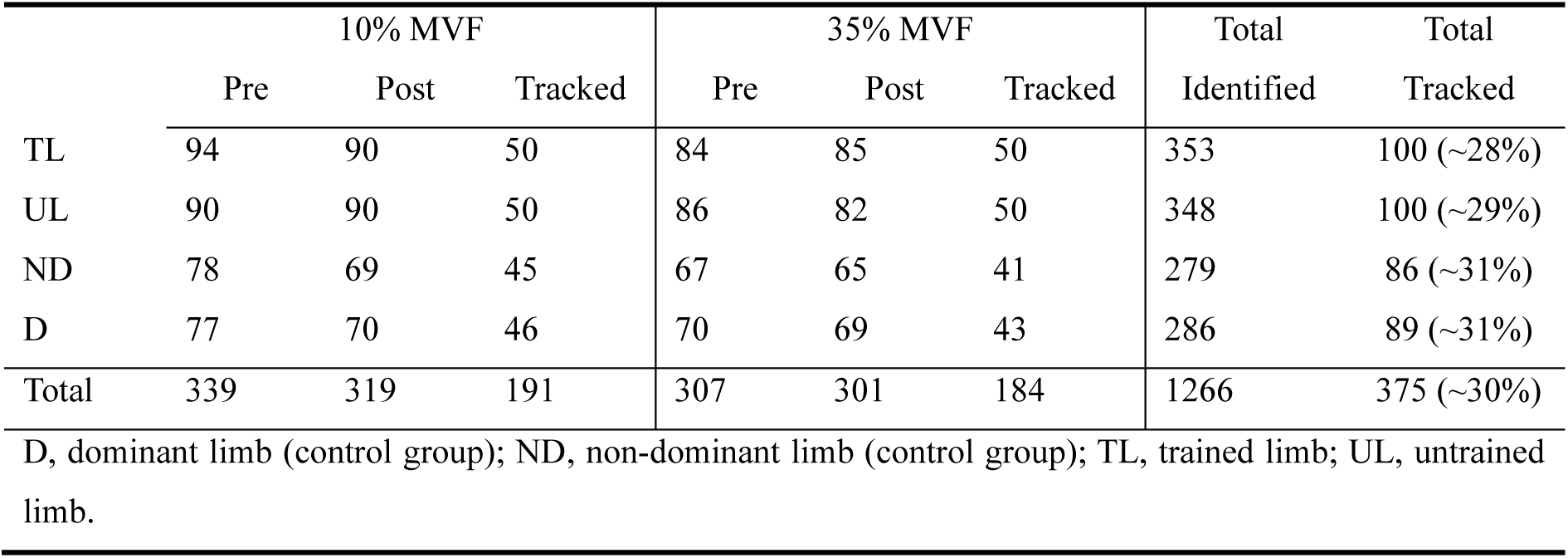
Identified and tracked motor units.

|  | 10% MVF |  |  | 35% MVF |  |  | Total | Total |
| --- | --- | --- | --- | --- | --- | --- | --- | --- |
|  | Pre | Post | Tracked | Pre | Post | Tracked | Identified | Tracked |
| TL | 94 | 90 | 50 | 84 | 85 | 50 | 353 | 100 (~28%) |
| UL | 90 | 90 | 50 | 86 | 82 | 50 | 348 | 100 (~29%) |
| ND | 78 | 69 | 45 | 67 | 65 | 41 | 279 | 86 (~31%) |
| D | 77 | 70 | 46 | 70 | 69 | 43 | 286 | 89 (~31%) |
| Total | 339 | 319 | 191 | 307 | 301 | 184 | 1266 | 375 (~30%) |

D, dominant limb (control group); ND, non-dominant limb (control group); TL, trained limb; UL, untrained limb.

The ICC values for comparisons of motor unit properties are presented in Table 3. All comparisons showed excellent test-retest reliability, reflecting the robustness of the employed methods and tracking procedures.

**Table 3:**
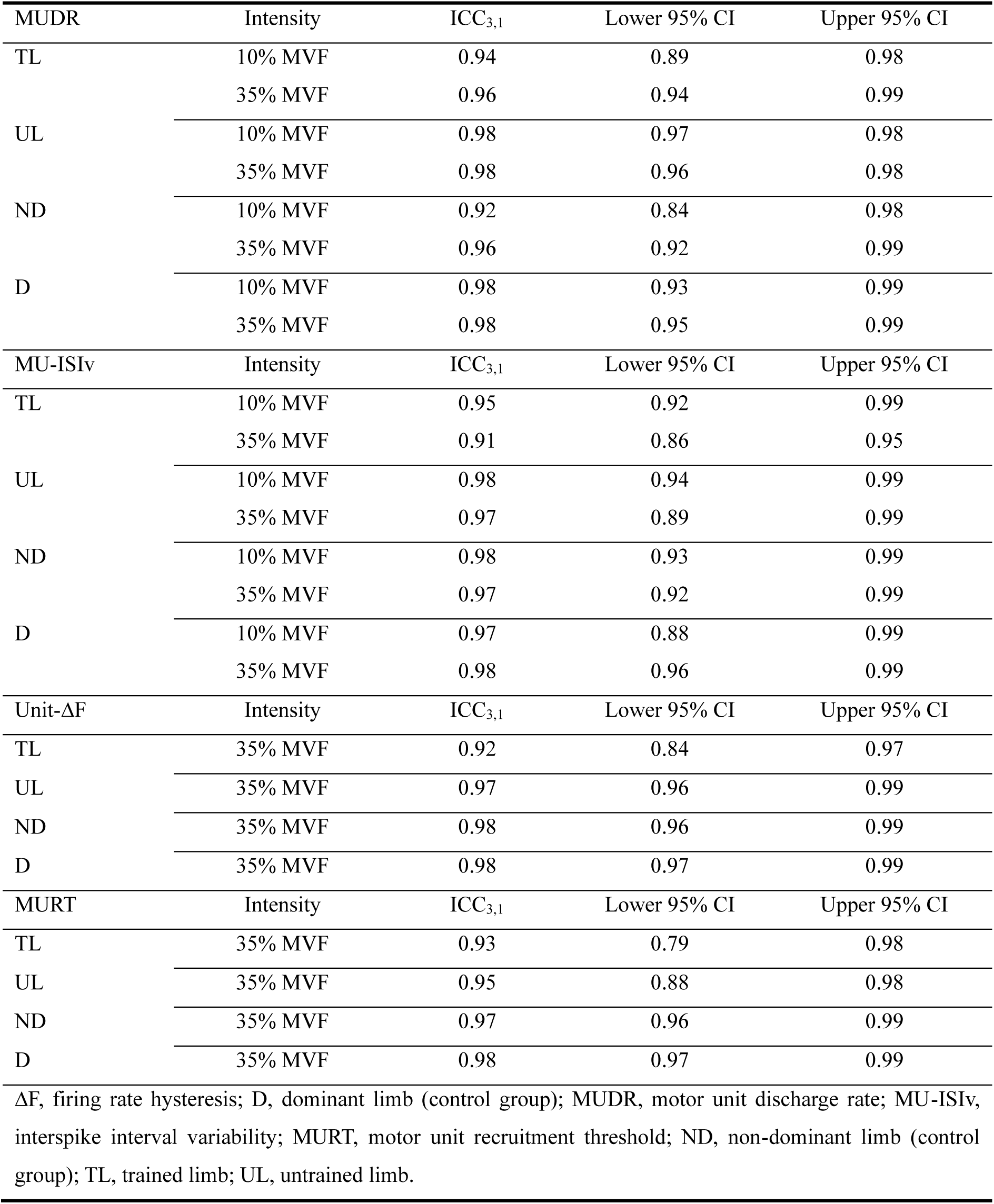
Test-retest reliability of tracked motor unit properties.

| MUDR | Intensity | ICC<sub>3,1</sub> | Lower 95% CI | Upper 95% CI |
| --- | --- | --- | --- | --- |
| TL | 10% MVF | 0.94 | 0.89 | 0.98 |
|  | 35% MVF | 0.96 | 0.94 | 0.99 |
| UL | 10% MVF | 0.98 | 0.97 | 0.98 |
|  | 35% MVF | 0.98 | 0.96 | 0.98 |
| ND | 10% MVF | 0.92 | 0.84 | 0.98 |
|  | 35% MVF | 0.96 | 0.92 | 0.99 |
| D | 10% MVF | 0.98 | 0.93 | 0.99 |
|  | 35% MVF | 0.98 | 0.95 | 0.99 |
| MU-ISI<sub>v</sub> | Intensity | ICC<sub>3,1</sub> | Lower 95% CI | Upper 95% CI |
| TL | 10% MVF | 0.95 | 0.92 | 0.99 |
|  | 35% MVF | 0.91 | 0.86 | 0.95 |
| UL | 10% MVF | 0.98 | 0.94 | 0.99 |
|  | 35% MVF | 0.97 | 0.89 | 0.99 |
| ND | 10% MVF | 0.98 | 0.93 | 0.99 |
|  | 35% MVF | 0.97 | 0.92 | 0.99 |
| D | 10% MVF | 0.97 | 0.88 | 0.99 |
|  | 35% MVF | 0.98 | 0.96 | 0.99 |
| Unit-ΔF | Intensity | ICC<sub>3,1</sub> | Lower 95% CI | Upper 95% CI |
| TL | 35% MVF | 0.92 | 0.84 | 0.97 |
| UL | 35% MVF | 0.97 | 0.96 | 0.99 |
| ND | 35% MVF | 0.98 | 0.96 | 0.99 |
| D | 35% MVF | 0.98 | 0.97 | 0.99 |
| MURT | Intensity | ICC<sub>3,1</sub> | Lower 95% CI | Upper 95% CI |
| TL | 35% MVF | 0.93 | 0.79 | 0.98 |
| UL | 35% MVF | 0.95 | 0.88 | 0.98 |
| ND | 35% MVF | 0.97 | 0.96 | 0.99 |
| D | 35% MVF | 0.98 | 0.97 | 0.99 |

ΔF, firing rate hysteresis; D, dominant limb (control group); MUDR, motor unit discharge rate; MU-ISI<sub>v</sub>, interspike interval variability; MURT, motor unit recruitment threshold; ND, non-dominant limb (control group); TL, trained limb; UL, untrained limb.

### Motor unit discharge rate, interspike interval variability, and recruitment thresholds

A significant *group x limb x time* interaction was found for MUDR at both 10% MVF (F_1, 339.15_ = 7.49, *p* = 0.007) and 35% MVF (F_1, 334.54_ = 42.07, *p* < 0.001). *Post hoc* analyses indicated a significant increase in MUDR following resistance training exclusively in trained limbs, at 10% MVF (ΔMUDR = + 2.43 pps [1.39, 3.46], *p* < 0.001, *d* = 0.66 [0.36, 0.97], Fig. 2A) and 35% MVF (ΔMUDR = + 2.45 pps [2.15, 2.74], *p* < 0.001, *d* = 2.24 [1.74, 2.73], Fig. 2B). Averaged results confirmed the linear mixed models, showing a significant MUDR increase only in trained limbs at 10%MVF (ΔMUDR = + 2.33 pps [1.63, 3.02], *p* < 0.001, *d* = 2.40 [1.13, 3.64], Fig. 2C) and 35% MVF (ΔMUDR = + 2.46 pps [2.02, 2.91], *p* < 0.001, *d* = 3.96 [2.05, 5.85], Fig. 2D). No significant differences were observed in the untrained limbs or the control group (*p* > 0.05).

**Fig. 2.**
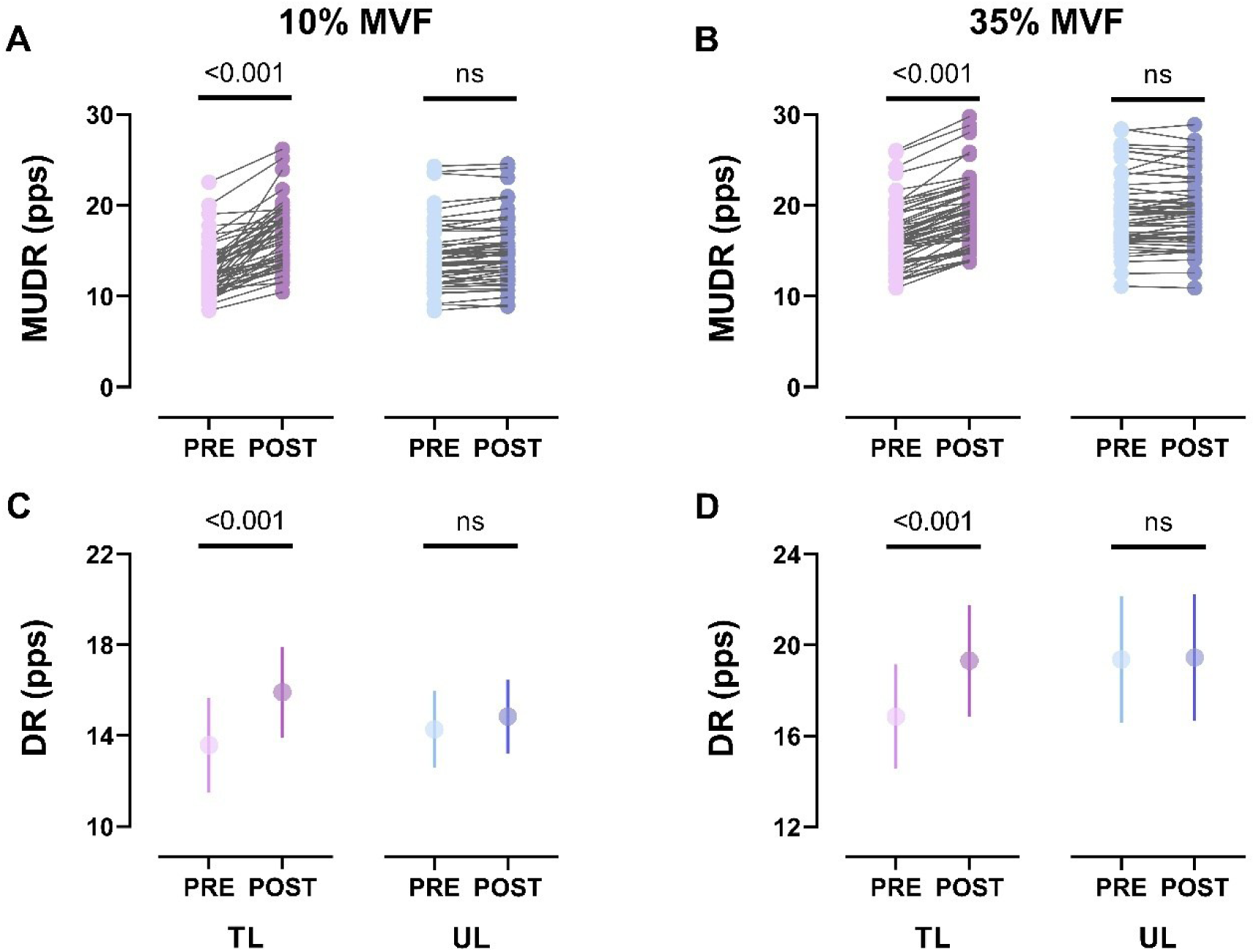
Intervention group MUDR and DR. MUDR comparisons for both limbs at 10% MVF (A) and 35% MVF (B) are displayed. Each marker represents an individual motor unit before and after the intervention. Averaged MUDR values for each participant are displayed as group-level comparisons at 10% MVF (C) and 35% MVF (D), represented as the mean with 95% CI. All results pertain to longitudinally tracked motor units.

A significant *group x time* interaction was observed for MU-ISIv at 10% MVF (F_1, 339.10_ = 7.74, *p* = 0.006) and 35% MVF (F_1, 40.75_ = 10.19, *p* = 0.003). *Post hoc* analyses revealed a significant decrease in MU-ISIv for both limbs at 10% MVF (trained, ΔMU-ISIv = - 2.18 % [-2.57, -1.78], *p* < 0.001, *d* = 1.57 [1.15, 1.98]; untrained, ΔMU-ISIv = - 1.77 % [- 2.08, - 1.45], *p* < 0.001, *d* = 1.58 [1.15, 1.99], Fig. 3A) and 35% MVF (trained, ΔMU-ISIv = - 3.51 % [- 4.18, - 2.85], *p* < 0.001, *d* = 1.43 [1.05, 1.81]; untrained, ΔMU-ISIv = - 2.87 % [- 3.35, - 2.39], *p* < 0.001, *d* = 1.71 [1.27, 2.15], Fig. 3B). Averaged results further confirmed a significant decrease in ISIv at both 10% MVF (trained, ΔISIv = - 2.09 % [-2.69, -1.49], *p* < 0.001, *d* = 2.49 [1.19, 3.76]; untrained, ΔISIv = - 1.76 % [- 2.21, - 1.31, *p* < 0.001, *d* = 2.80 [1.37, 4.20], Fig. 3C) and 35% MVF (trained, ΔISIv = - 3.72 % [-5.01, -2.43], *p* < 0.001, *d* = 2.49 [1.19, 3.76]; untrained, ΔISIv = - 2.86 % [- 3.74, - 1.98, *p* < 0.001, *d* = 2.33 [1.09, 3.54], Fig. 3D). No significant differences were observed in the CNT group (*p* > 0.05).

**Fig 3.**
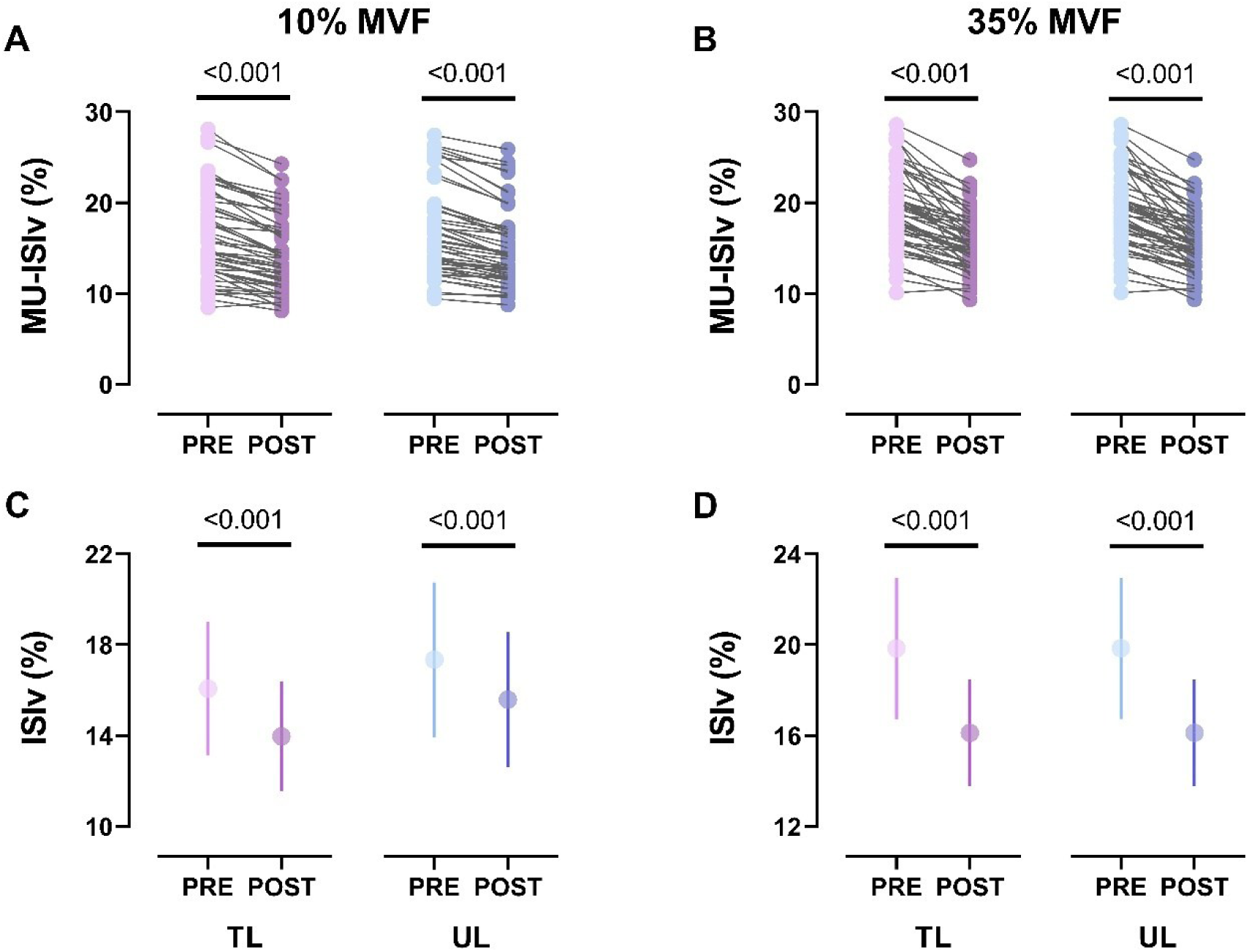
Intervention group MU-ISIv and ISIv. MU-ISIv comparisons for both limbs at 10% MVF (A) and 35% MVF (B) are shown, with each marker representing an individual motor unit before and after the intervention. Averaged ISIv values are presented as group-level comparisons at 10% MVF (C) and 35% MVF (D), depicted as the mean with 95% CI. All results pertain to longitudinally tracked motor units.

A significant *group x limb x time* interaction was found for absolute MURT (F_1, 334.28_ = 15.64, *p* < 0.001, Fig. 4 A-C). *Post hoc* analyses revealed a significant decrease in absolute MURT for both limbs (trained, ΔMURT = - 6 N [-8, -4], *p* < 0.001, *d* = 1.01 [0.68, 1.33]; untrained, ΔMURT = - 3 N [- 4, - 2], *p* < 0.001, *d* = 1.23 [0.86, 1.60]). Averaged results confirmed a significant decrease in absolute RT (trained, ΔRT = - 6 N [-9, -3], *p* < 0.001, *d* = 1.53 [0.58, 2.45]; untrained, ΔRT = - 3 N [- 5, - 1], *p* < 0.001, *d* = 1.70 [0.69, 2.68]). A significant *group x time* interaction was also observed in the relative MURT (F_1, 341.53_ = 38.61, *p* < 0.001, Fig. 4 B-D). *Post hoc* analyses revealed a significant decrease in relative MURT for both limbs (trained, ΔMURT = - 4.9 % [-5.5, -4.3], *p* < 0.001, *d* = 2.24 [1.74, 2.73]; untrained, ΔMURT = - 2.7 % [- 3.0, - 2.3], *p* < 0.001, *d* = 2.29 [1.75, 2.81]). Averaged results confirmed a significant decrease in absolute RT (trained, ΔRT = - 5.2 % [-6.4, -3.9], *p* < 0.001, *d* = 3.00 [1.49, 4.48]; untrained, ΔRT = - 2.7 % [- 3.3, - 2.0], *p* < 0.001, *d* = 2.82 [1.38, 4.22]). No significant differences were observed in the control group for both absolute and relative recruitment threshold forces (*p* > 0.05).

**Fig 4.**
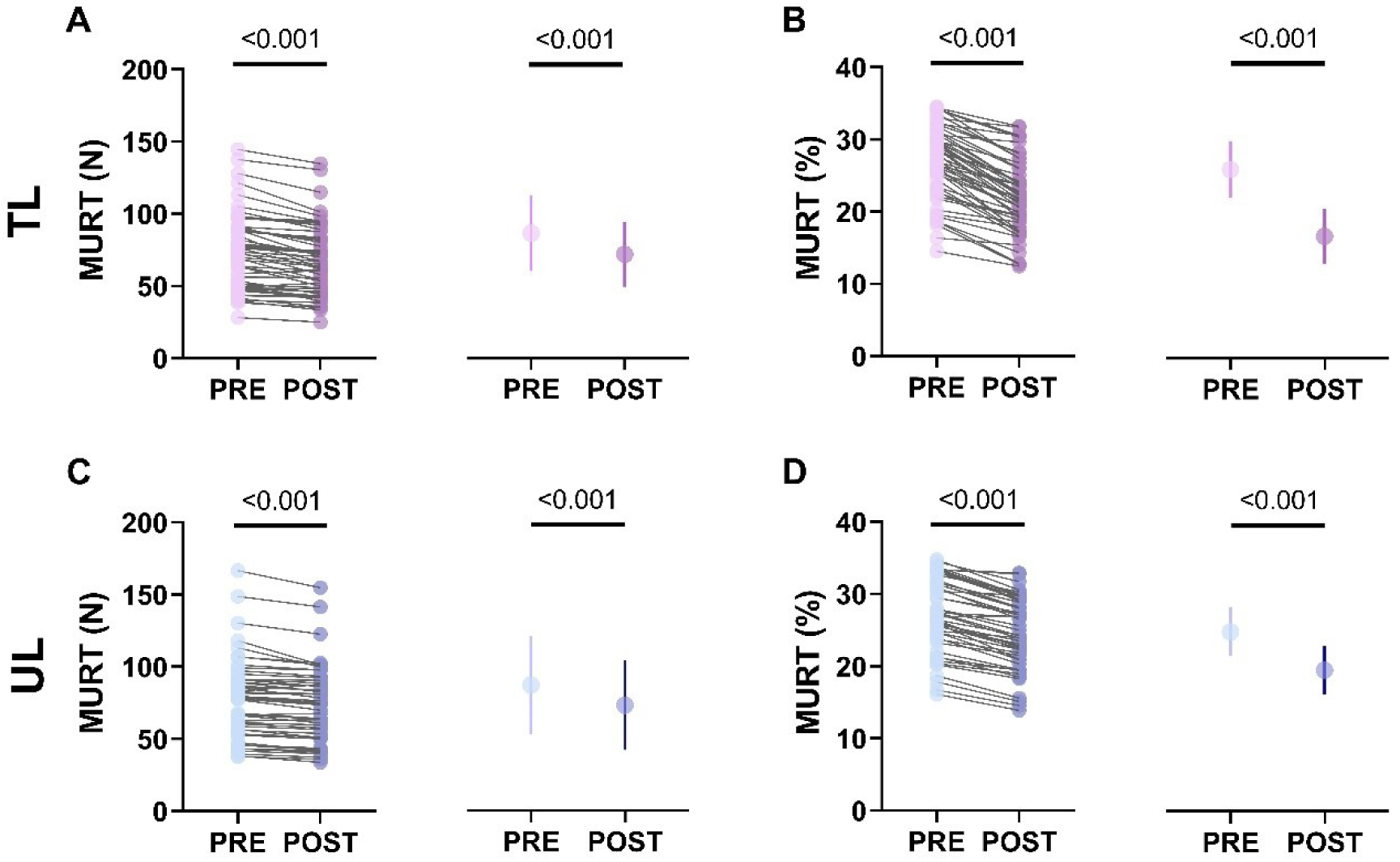
Intervention group MURT and RT in 35% MVF contractions. Absolute (A) and relative (B) MURT for trained limbs, as well as absolute (C) and relative (D) MURT for untrained limbs are displayed, with each marker representing an individual motor unit before and after the intervention. Averaged RT are reported as the mean with 95%CI. All results pertain to longitudinally tracked motor units.

### Coherence and CSI-V

At 10% MVF, pairwise comparisons revealed a significantly higher mean z-coherence in the δ bandwidth for both the trained (ΔZC = + 0.405 [0.166, 0.644], *p* = 0.004, *d* = 1.21 [0.36, 2.02], Fig. 5A) and the untrained limbs (ΔZC = + 0.619 [0.327, 0.912], *p* < 0.001, *d* = 1.52, [0.57, 2.43], Fig. 5C). No significant differences were observed for α and β bandwidths (*p* > 0.05). Additionally, the maximal z-coherence increased in the δ bandwidth for both trained (ΔZC = + 0.698 [0.355, 1.040], *p* = 0.001, *d* =1.46 [0.53, 2.35]) and untrained limbs (ΔZC = + 0.658 [0.350, 0.967], *p* < 0.001, *d* = 1.53 [0.58, 2.44]). At 35% MVF, a significant increase in the mean z-coherence in the δ bandwidth was observed for both the trained (ΔZC = + 0.800 [0.487, 1.114], *p* < 0.001, *d* = 1.83 [0.77, 2.84], Fig. 5B) and the untrained limbs (ΔZC = + 0.524 [0.303, 0.744], *p* < 0.001, *d* =1.70 [0.69, 2.67], Fig. 5D). No significant differences were observed for α and β bandwidths (*p* > 0.05). The maximal z-coherence increased in the δ bandwidth for both the trained (ΔZC = + 0.871 [0.646, 1.097], *p* < 0.001, *d* = 2.76 [1.35, 4.15]) and the untrained limbs (ΔZC = + 0.513 [0.247, 0.779], *p* = 0.002, *d* = 1.38 [0.48, 2.24]). No significant differences were observed in the control group across all bandwidths (*p* > 0.05). Mean coherence results are reported in Table 4.

**Fig. 5.**
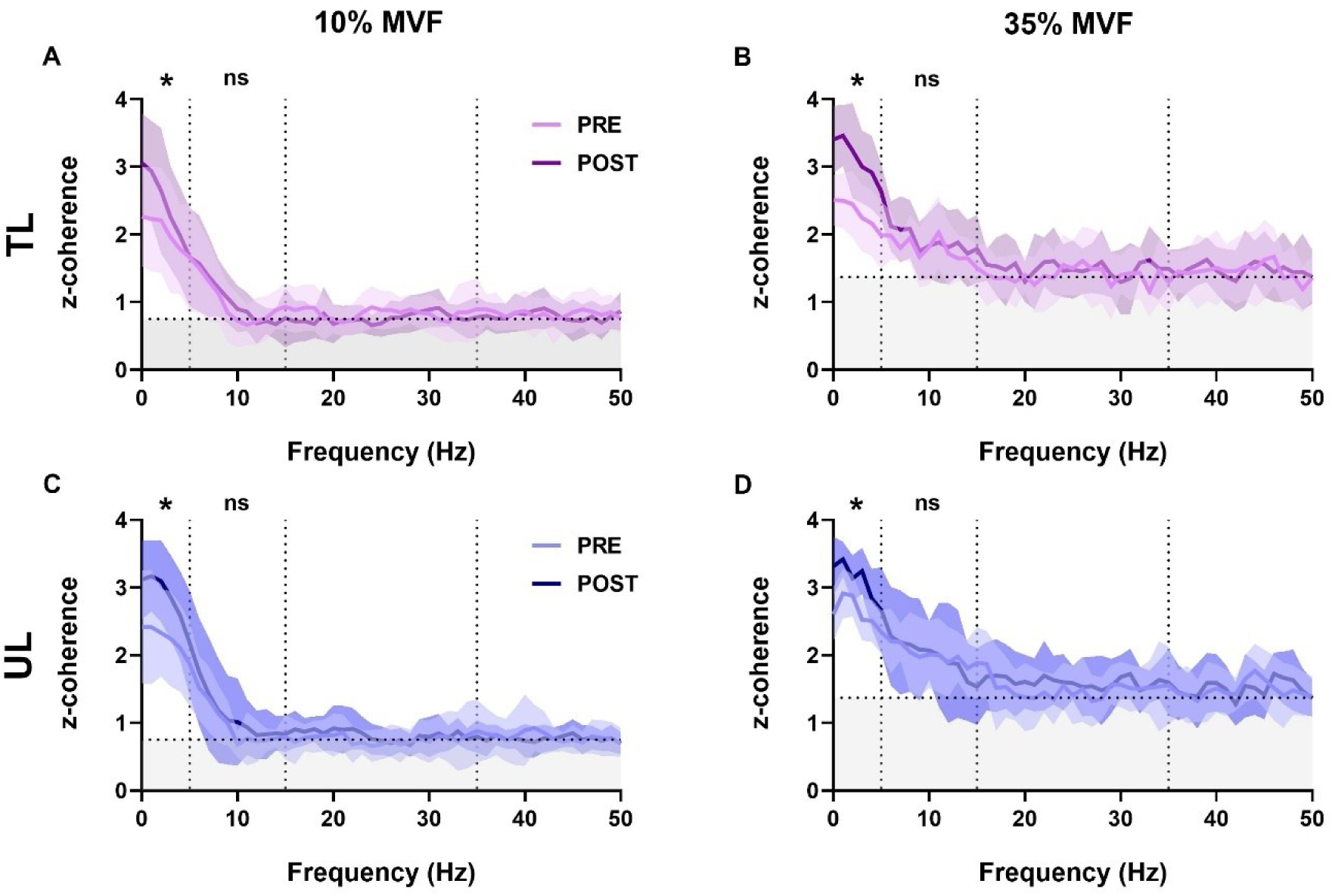
Mean z-coherence profile comparisons of trained and untrained limbs. Z-coherence comparisons for trained limbs at 10% MVF (A) and 35% MVF (B) are shown, alongside untrained limb z-coherence at 10% MVF (C) and 35% MVF (D). All coherence profiles are derived from longitudinally tracked motor units. The solid line represents the mean z-coherence across participants, while the shaded area indicates the SD. The horizontal dotted line marks the significant coherence threshold, identified as the 95th percentile (0.83 at 10% MVF and 1.37 at 35% MVF). Vertical dotted lines delineate the (0 - 5 Hz) δ and (5 - 15 Hz) α. Significant comparisons within specific bandwidths are denoted by the symbol ‘*’.’

**Table 4:** mean z-coherence.

| MVF | INT |  |  |  | CNT |  |  |  |
| --- | --- | --- | --- | --- | --- | --- | --- | --- |
|  | TL |  | UL |  | ND |  | D |  |
| 10% | Pre | Post | Pre | Post | Pre | Post | Pre | Post |
| $\delta$ | 2.02±0.7 | 2.43±0.7 * | 2.23±0.7 | 2.85±0.6 * | 2.06±0.5 | 2.11±0.5 | 2.19±0.6 | 2.28±0.6 |
| $\alpha$ | 0.95±0.3 | 0.98±0.3 | 0.95±0.2 | 1.15±0.4 | 1.24±0.3 | 1.27±0.3 | 1.15±0.3 | 1.22±0.4 |
| 35% | Pre | Post | Pre | Post | Pre | Post | Pre | Post |
| $\delta$ | 2.31±0.4 | 3.11±0.5 * | 2.63±0.4 | 3.15±0.4 * | 2.32±0.6 | 2.34±0.6 | 2.54±0.6 | 2.49±0.7 |
| $\alpha$ | 1.78±0.3 | 1.99±0.3 | 2.00±0.3 | 2.10±0.3 | 1.58±0.5 | 1.69±0.5 | 1.47±0.3 | 1.60±0.4 |

D, dominant limb (control group); ND, non-dominant limb (control group); TL, trained limb; UL, untrained limb. Paired t-tests were used for statistical comparisons. Data are presented as the mean±SD. Significant pairwise interactions are indicated with the symbol ‘\*’. B results are reported for information completeness.

At 10% MVF, pairwise comparisons revealed a significantly lower CSI-V for both the trained (ΔCSI-V = - 0.22 [-0.36, -0.08], *p* = 0.006, *d* = 1.12 [0.30, 1.91]) and untrained limbs (ΔCSI- V = - 0.15 [-0.24, -0.06], *p* = 0.004, *d* = 1.19 [0.35, 2.00]). Similarly, decreased CSI-V was also observed at 35% MVF in both the trained (ΔCSI-V = - 0.33 [-0.40, -0.26], *p* < 0.001, *d* = 3.46 [1.76, 5.13]) and untrained sides (ΔCSI-V = - 0.21 [-0.27, -0.15], *p* < 0.001, *d* = 2.37 [1.12, 3.60]).

### Firing rate hysteresis

We identified a total of 196 unit-wise observations, including baseline and post-intervention 35% MVF trapezoidal contractions. This number corresponds to 98 longitudinally tracked motor units across the intervention: 29 from trained limbs, 25 from untrained limbs, 22 from dominant limbs, and 22 from non-dominant limbs. A significant *group x limb x time* interaction was observed for the unit-ΔF (F_1, 170.62_ = 4.82, *p* = 0.008). *Post hoc* analyses revealed a significant increase for the trained limbs following the 4-week intervention (unit-ΔF = + 1.61 pps [1.22, 2.00], *p* < 0.001, *d* = 1.58 [1.02, 2.12], Fig. 6A). No significant differences were observed for untrained limbs or the control group (*p* > 0.05, Fig. 6D). Averaged comparisons also demonstrated a significant increase for the trained limbs (participant-ΔF = + 1.65 pps [1.12, 2.17], *p* < 0.001, *d* = 2.26 [1.04, 3.44]), with no differences for untrained limbs or the control group (*p* > 0.05).

**Fig. 6.**
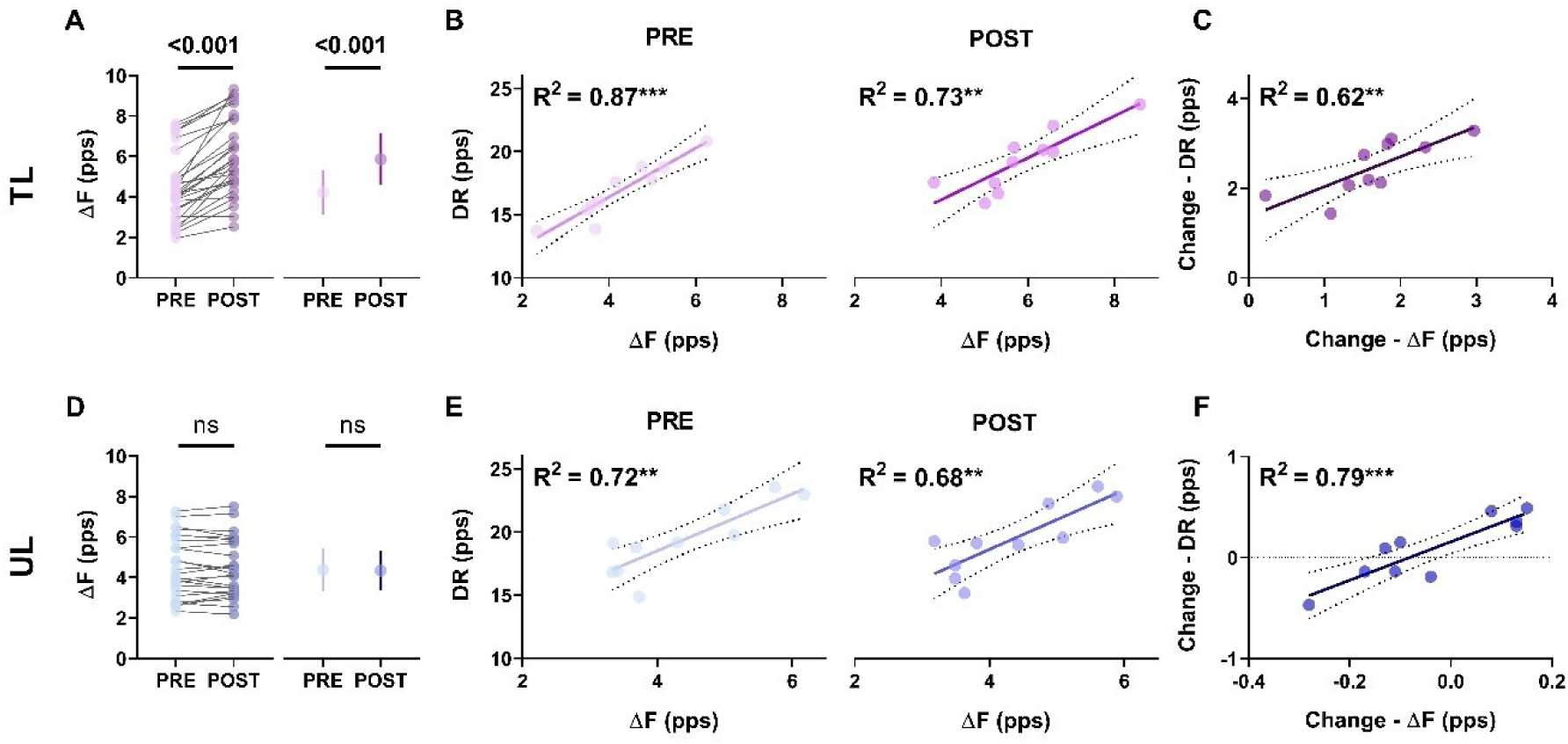
Firing rate hysteresis and the gain in DR at 35% MVF contractions. Comparisons of unit-ΔF and participant-ΔF in trained (A) and untrained (D) limbs show a significant increase in the estimates of the persistent inward currents only on the trained side. The associations between the averaged DR as a function of ΔF are presented for trained (B) and untrained (E) limbs, with the relative association between the change in DR as a function of the change in ΔF for the trained (C) and untrained (F) sides. All data refer to longitudinally tracked motor units. Averaged results are presented as the mean and 95% CI. ** *p* < 0.01; *** *p* < 0.001. TL, trained limbs; UL, untrained limbs.

The association between ΔF and DR was very strong at both baseline and post-intervention for the trained (PRE, R^2^ = 0.87, *p* < 0.001; POST, R^2^ = 0.73, *p* = 0.001, Fig. 6B) and untrained limbs (PRE, R^2^ = 0.72, *p* = 0.002; POST, R^2^ = 0.68, *p* = 0.003, Fig. 6E). A strong association was also observed between the absolute change in DR and the absolute change in ΔF for both the trained (R^2^ = 0.62, *p* = 0.006, Fig. 6C) and untrained sides (R^2^ = 0.79, *p* < 0.001, Fig. 6F). These results indicate a robust relationship between the PIC-related contribution to intrinsic motoneuron properties and their neural output to muscles, with changes in one variable strongly mirroring changes in the other in both limbs. Therefore, significant modifications in trained side ΔF correlate with significant differences in DR, while fluctuations in untrained limb ΔF were strongly associated with DR values, not altered with training.

### Associations between neural and force variables

In the trained limbs, the change in MVF was strongly associated with the change in all the neural variables (δ z-coherence, R^2^ = 0.74, *p* = 0.001; RT, R^2^ = 0.70, *p* = 0.003; ΔF, R^2^ = 0.85, *p* < 0.001; DR, R^2^ = 0.84, *p* < 0.001; Fig. 7A). In contrast, in the untrained limbs the change in MVF was significantly associated only with δ z-coherence (R^2^ = 0.75, *p* = 0.001) and RT (R^2^ = 0.66, *p* = 0.004; Fig. 8B), with no significant associations observed with the change in ΔF (R^2^ = 0.15, *p* = 0.268) and DR (R^2^ = 0.17, *p* = 0.222). These results suggest that increased muscle strength in trained limbs was associated with enhanced relative shared synaptic input, earlier motoneuron activation, broader PIC estimates, and higher neural drive. In contrast, the increase in untrained limb MVF was exclusively associated with a higher proportion of common synaptic input and earlier motoneuron activation, with no contribution from PIC amplitude or neural drive.

**Fig. 7.**
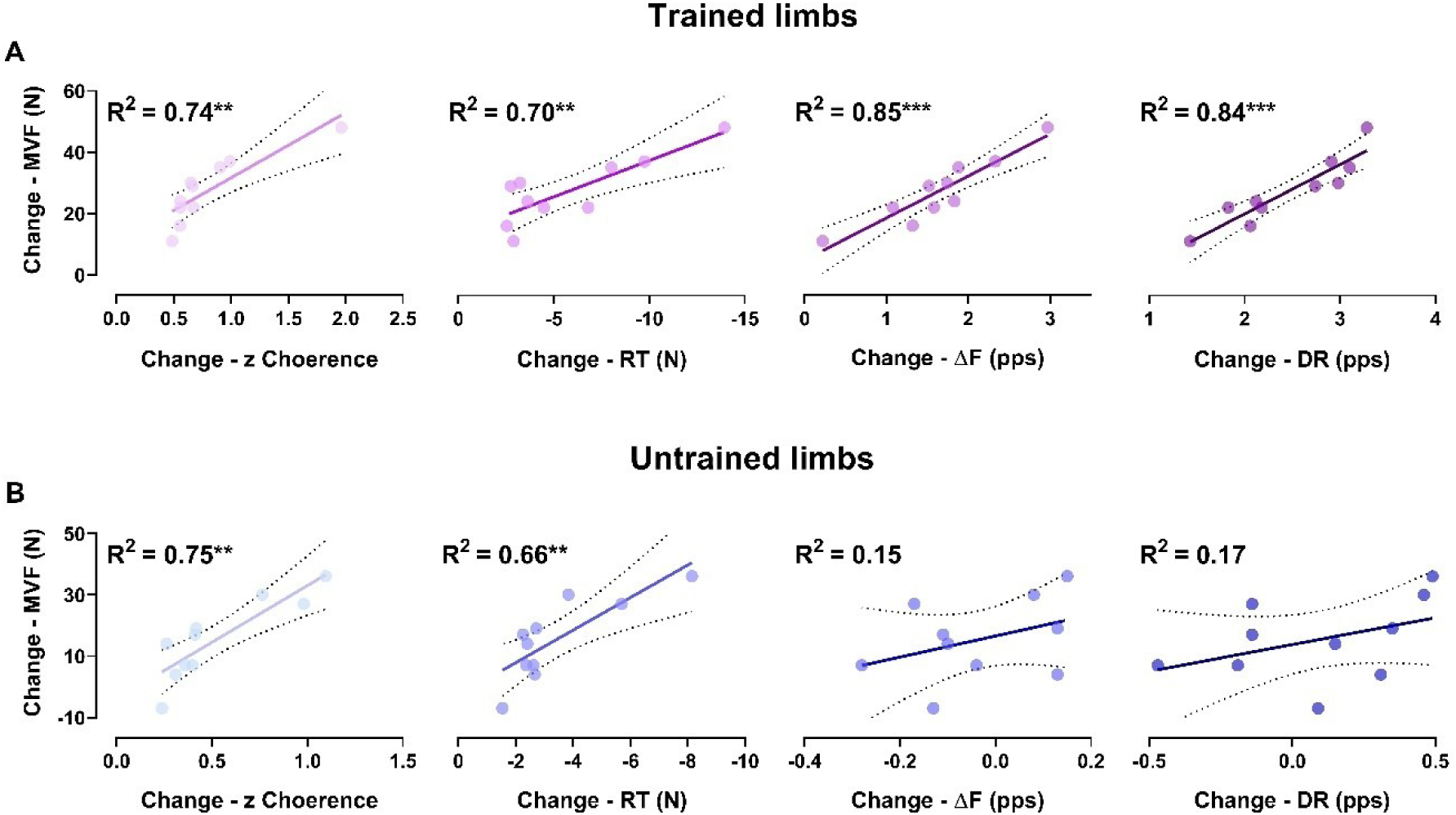
Change in MVF as a function of the change in neural variables. For trained limbs (A) and untrained limbs (B), the associations between the change in MVF as a function of the change in the neural parameters are presented for trapezoidal contractions at 35% MVF results. Each marker refers to a participant’s value. ** *p* < 0.01; *** *p* < 0.001. TL, trained limbs; UL, untrained limbs.

**Fig. 8.**
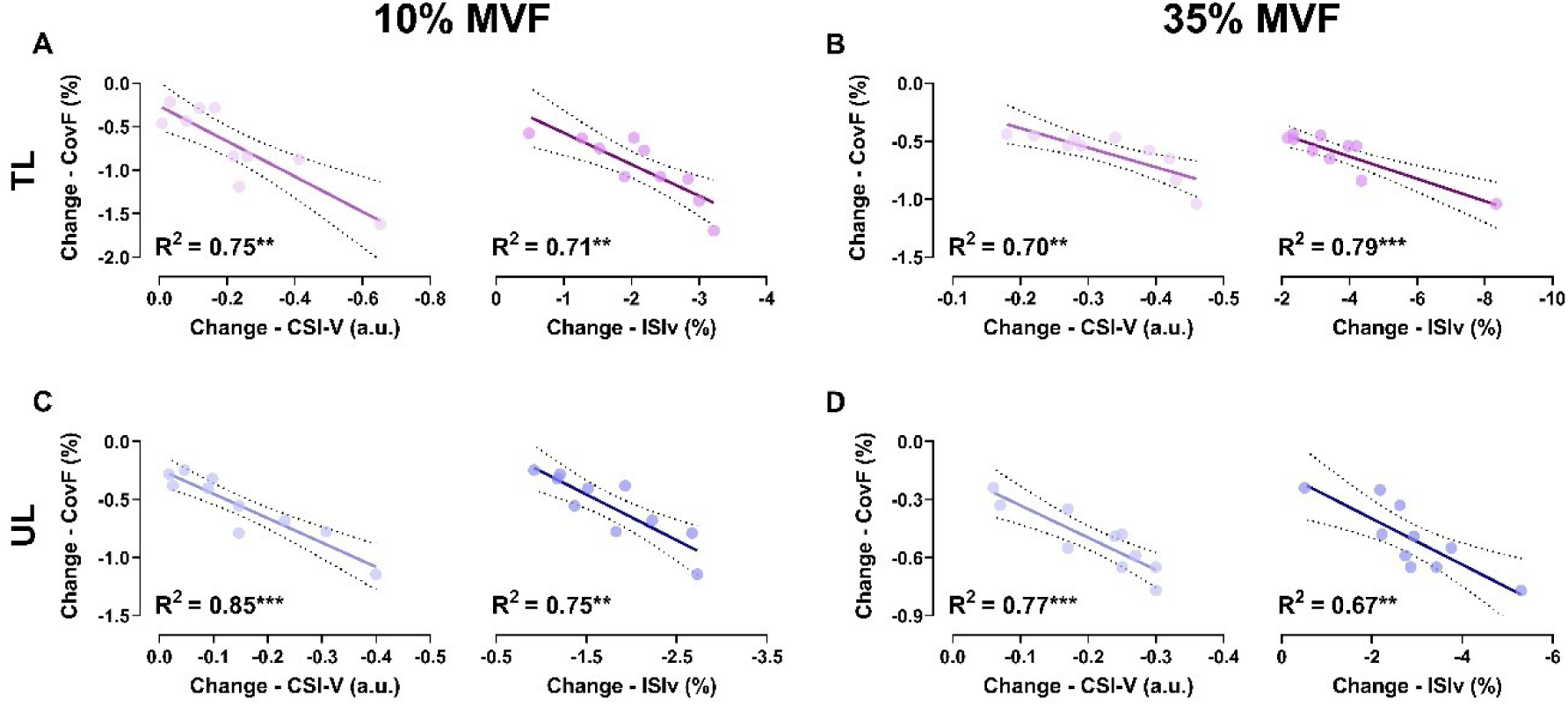
Change in CovF as a function of the change in neural variables. Scatter plots display the association between the change in CovF as a function of CSI-V and ISIv at 10% MVF and 35% MVF in trained (A-B) and untrained limbs (C-D). Each marker refers to a participant’s value. ** *p* < 0.01; *** *p* < 0.001. TL, trained limbs; UL, untrained limbs.

Enhanced force steadiness, reflected by reduced CovF, was strongly associated with the change in all the neural variables in trained limbs at 10% MVF (CSI-V, R^2^ = 0.75, *p* = 0.001; ISIv, R^2^ = 0.71, *p* = 0.002; Fig. 8A) and 35% MVF (CSI-V, R^2^ = 0.70, *p* = 0.003; ISIv, R^2^ = 0.79, *p* < 0.001; Fig. 8B). Similarily, decreased CovF was strongly associated with neural variable adaptations in untrained limbs at both 10% MVF (CSI-V, R^2^ = 0.85, *p* < 0.001; ISIv, R^2^ = 0.75, *p* = 0.001; Fig. 8C) and 35% MVF (CSI-V, R^2^ = 0.77, *p* < 0.001; ISIv, R^2^ = 0.67, *p* = 0.004; Fig. 8D). These findings highlight a parallel increase in neural variables associated with force accuracy with the steadiness in the mechanical output in both limbs.

### Parallel neuromechanical adaptations in trained and untrained limbs

Concerning neuromechanical variables associated with absolute force production, both trained and untrained limbs exhibited parallel adaptations, with strong associations in δ z-coherence (R^2^ = 0.74, *p* = 0.001), RT (R^2^ = 0.85, *p* < 0.001), and MVF (R^2^ = 0.90, *p* < 0.001, Fig. 9A). Similarly, parallel adaptations in variables related to force steadiness were observed, with robust associations in CSI-V (R^2^ = 0.81, *p* < 0.001), ISIv (R^2^ = 0.83, *p* < 0.001), and CovF (R^2^ = 0.77, *p* < 0.001, Fig. 9B). These results do not imply that the magnitude of responses is identical between trained and untrained limbs; rather, our findings suggest that the observed neuromechanical adaptations are proportional on both sides.

**Fig. 9.**
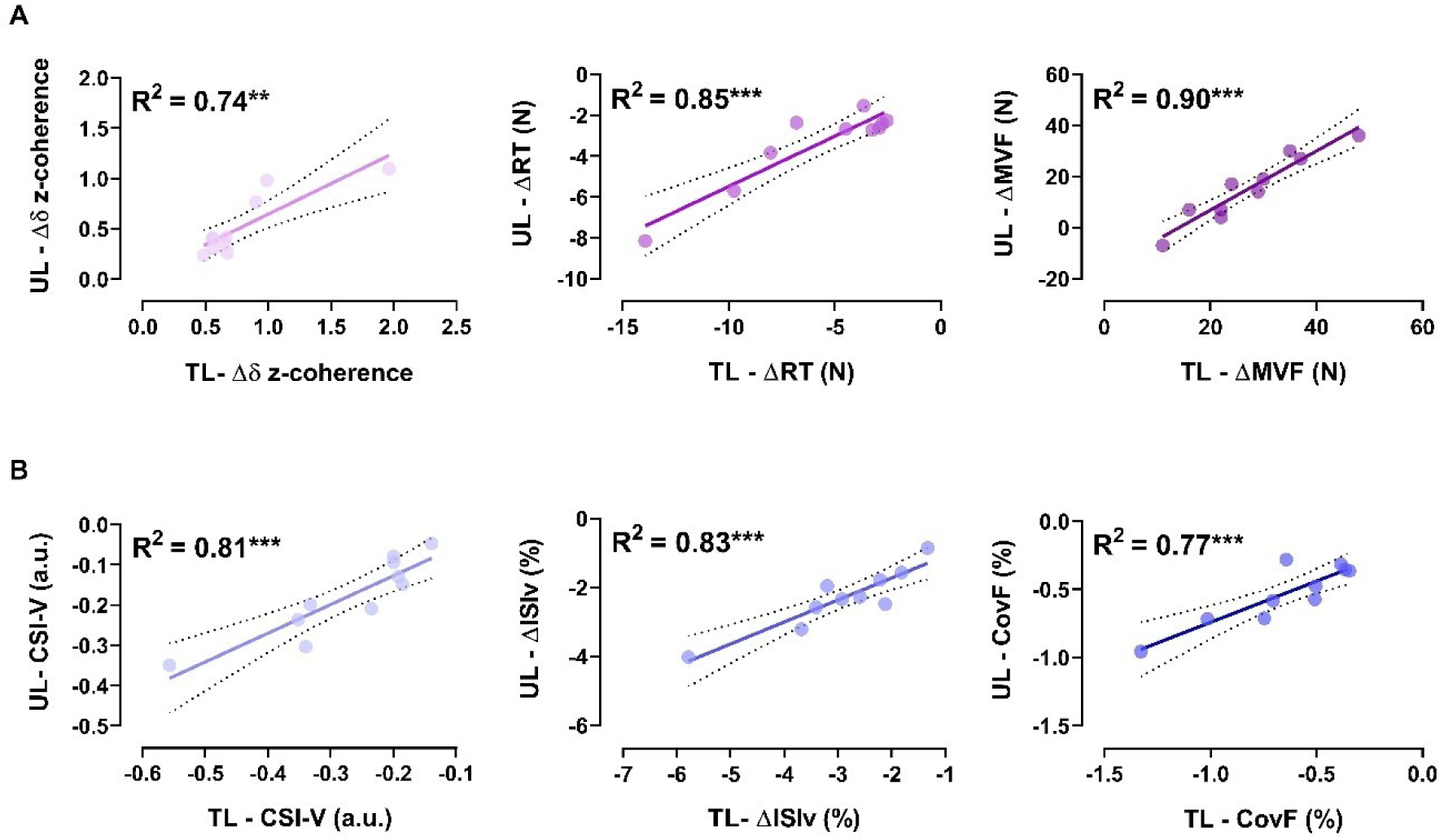
Association between untrained – trained limbs adaptations. Scatter plots illustrate the associations between neuromechanical responses in untrained - trained limbs. Panel A depicts adaptations related to improved force production, while Panel B shows those related to enhanced force steadiness. In Panel B, data represent the average responses recorded between 10% and 35% MVF. Each marker refers to a participant’s value. ** *p* < 0.01; *** *p* < 0.001. TL, trained limbs; UL, untrained limbs.

## Discussion

A 4-week unilateral eccentric resistance training elicited significant neuromechanical adaptations in both trained and untrained muscles. On the exercised limb, strength gains were accompanied by an increased proportion of common synaptic input, earlier motor unit activation, greater estimated PIC amplitude, and higher motor unit discharge rates. In contrast, the enhanced muscle strength observed in the contralateral untrained limbs was associated with an increased proportion of shared synaptic inputs and reduced motor unit recruitment thresholds without alterations in PIC amplitude. Concurrently, force variability decreased together with the variance in common input and the fluctuation in spiking activity in both limbs. Notably, adaptations on the untrained side were proportional to those on the trained side, suggesting that gain transfers are plausibly mediated by adaptations in shared input and a suppressed synaptic noise. Additional strength improvements in trained limbs are plausibly attributable to enhanced intrinsic motoneuron properties and neural drive.

### Methodological considerations, accuracy, and reliability

The number of identified and tracked motor units in the present study was consistent with expectations based on HDsEMG decomposition from biceps brachii, as reported in previous studies (Casolo et al., 2021; Del Vecchio, Holobar, et al., 2020; Duchateau & Enoka, 2022; Frančič & Holobar, 2022). Longitudinal tracking enabled the identification of the same motor units at baseline and after resistance training, addressing the limitation of sampling different motor units at each time point. This methodological approach provides valuable insights into the neural mechanisms underpinning cross-education at the motor unit level (Del Vecchio, Casolo, et al., 2019; Martinez-Valdes et al., 2017). Approximately 30% of the identified motor units were successfully tracked, aligning with previous findings involving resistance training and HDsEMG acquisition (Del Vecchio, Casolo, et al., 2019). The excellent test-retest reliability observed across groups and sides supports the accuracy of the tracking procedures, reinforcing the robustness of the results (Goodlich, Del Vecchio, & Kavanagh, 2023; Goodlich, Del Vecchio, Horan, et al., 2023). Notably, high ICC observed for estimates of persistent inward currents further confirms the reliability of the paired motor unit approach for non-invasive assessment of PIC contributions to intrinsic motoneuron properties during voluntary ramp contractions (Goodlich, Del Vecchio, Horan, et al., 2023; Mesquita et al., 2023, 2024; Orssatto et al., 2021; Wilson et al., 2015), demonstrating sensitivity in detecting the contribution of PICs to motoneuron properties in response to different interventions (Goodlich et al., 2024; Orssatto et al., 2023).

### Neural adaptations mediating increased trained muscle strength

In the present study, resistance training significantly increased muscle strength and skill on the trained side, supporting previous evidence of neural adaptations mediating such responses (Škarabot et al., 2021). Our findings indicate that these adaptations include a higher proportion of relative shared synaptic input, lower motor unit recruitment thresholds, and increased discharge rates accompanied by elevated PIC amplitude, all of which are recognised contributors to enhanced muscle force (Del Vecchio, Casolo, et al., 2019; Del Vecchio et al., 2024; J. Dideriksen & Del Vecchio, 2023; Orssatto et al., 2023). Furthermore, the observed improvements in force steadiness were associated with a reduction in the variability of common synaptic input and decreased interspike variability, which translated into significantly lower force fluctuation during steady contraction phases (Enoka & Farina, 2021; Feeney et al., 2018; Vila-Chã & Falla, 2016).

### Neural mechanisms of cross-education

Enhancements of strength and skill in the contralateral untrained side following unilateral resistance training have been previously attributed to multiple neural mechanisms (Carson, 2005; Farthing et al., 2007; Fimland et al., 2009; Lecce, Conti, et al., 2025; Ruddy & Carson, 2013). Among these, we have recently demonstrated that cross-education at the motor unit level is associated with higher net discharge rates, lower motor unit recruitment thresholds, and reduced spiking variability (Lecce, Conti, et al., 2025). However, it was not possible to clarify the mechanisms associated with the relative shared synaptic input, its absolute variance, and the role of PICs in mediating these adaptations. In the present study, we extended these findings by reporting novel neural adaptations observed for the first time in the contralateral untrained limb. Specifically, we observed a greater relative shared synaptic input and reduced variability in low-frequency common oscillations without contribution from PICs. These adaptations were associated with lower motor unit recruitment thresholds, reduced variability in the spiking activity, higher maximal force output, and enhanced force steadiness.

### Shared synaptic input, synaptic noise, and mechanical output

Our findings revealed that both limbs exhibited higher relative shared input to motoneurons alongside a decrease in the recruitment threshold forces, with these neural responses being strongly associated with a higher absolute force production. These results support our previous hypotheses that earlier motoneuron activation resulted from modifications in shared synaptic inputs, thereby leading to increased absolute force output (J. Dideriksen & Del Vecchio, 2023; Lecce, Conti, et al., 2025). Accordingly, lower motor unit recruitment thresholds reflect the activation of a larger pool of motor units during force production, implying a greater mechanical output (J. Dideriksen & Del Vecchio, 2023; Škarabot et al., 2021). This reflects that more motor units are active at a relative force level without necessarily implying a greater strength of PIC amplitude contributing to intrinsic motoneuron properties (Kim et al., 2019). These adaptations would explain the increase in untrained muscle force without a relative increase in the neural drive, further supporting the contribution of enhanced motor unit recruitment in mediating cross-education (Fimland et al., 2009; Green & Gabriel, 2018a; Manca et al., 2018).

Concurrently, our findings indicate a reduction in the variance of common input within the low-frequency bandwidth (δ), accompanied by decreased variability in the spiking activity of motor units in both the exercised and contralateral motoneuron pools. Low-frequency oscillations are known to modulate force output, and a reduced variance in these correlated inputs translates into a more consistent neural drive, thereby reducing force variability (J. L. Dideriksen et al., 2012; Negro et al., 2009, 2016; Rodriguez-Falces et al., 2017). This is attributed to the motoneuron pool, which has been demonstrated to act as a low-pass filter that preserves the common synaptic input while attenuating high-frequency oscillations and independent components (Farina et al., 2014). Consequently, when common input fluctuations are suppressed, its near-undistorted transmission through the motoneurons implies a more stable neural drive (Negro & Farina, 2011). This mechanism accounts for our observations of lower fluctuations in the low-frequency components of motoneuron activity and a reduced interspike interval variability. Essentially, the decreased variability in the common synaptic input, combined with greater suppression of random membrane noise, results in a more regular motoneuron output to muscles. As the effective neural drive directly mirrors these (*correlated*) inputs, the reduced variance observed in response to resistance training is then reflected in improved muscle force control, leading to enhanced force steadiness (A. Castronovo et al., 2018; Enoka & Farina, 2021; Feeney et al., 2018).

Notably, low-frequency components mirror signals from cortical projections and spinal circuitries converging onto spinal motoneurons (Hug et al., 2023). Therefore, our findings likely stem from adaptations occurring upstream of the motoneurons, suggesting that supraspinal or spinal mechanisms may have contributed to motor unit adaptations, facilitating the transfer of strength and skill from the exercised limb to the contralateral limb. This interpretation aligns with existing evidence on cross-education (Carson, 2005, 2020; Frazer et al., 2018; Hendy & Lamon, 2017; Ruddy & Carson, 2013).

### PIC-amplitude and neural drive

Enhanced estimates of the persistent inward currents and MUDR were exclusively observed in the trained limb, with no significant changes detected in the untrained side. The strong association between PIC estimates and MUDR in both limbs underscores the strong association between PIC-related variable gain system and discharge rate modulation (Heckman et al., 2005; Orssatto et al., 2021). In absolute terms, significant increases in PIC amplitudes were accompanied by a significantly higher MUDR, reflecting an enhanced capacity of motoneurons to sustain elevated discharge rates on the trained side, while no significant changes were observed on the untrained side. These findings highlight the role of PICs in facilitating increased motoneuron discharge rates in response to repetitive mechanical overload, specifically within the exercised motoneuron pool (Del Vecchio, Casolo, et al., 2019; Orssatto et al., 2023). The precise mechanisms underlying the observed modifications in both PICs and MUDR following strength training interventions remain unclear. Although speculative, several potential mechanisms may account for these differential adaptations.

Evidence from animal studies indicates that resistance training induces physiological changes in motoneuron properties, such as shorter spike durations, increased input resistance, a lower rheobase, and a reduced minimum current required to initiate rhythmic firing (Krutki et al., 2015, 2017). These adaptations are also associated with increased maximum spike frequencies during early and steady-state motoneuron activity following resistance training (Dai et al., 2024). Furthermore, it has been observed that motor units in trained limbs generate greater muscle forces, with higher discharge rates at the same depolarisation current, inducing enhanced performance in response to intervention (Dai et al., 2024; Krutki et al., 2017). Such adaptations imply that mechanical overloading, or the absence thereof, may trigger physiological modifications in motoneurons, likely due to chronic alterations in ion channel conductances (Cormery et al., 2005).

Furthermore, it is plausible that the monoaminergic system would eventually influence these adaptations (Heckman et al., 2009; Mesquita et al., 2024). For instance, increased serotonin immunoreactivity in the hypoglossal nucleus has been observed following tongue resistance training in rats, suggesting an increased serotonergic input onto motoneurons innervating the target muscle fibres (Behan et al., 2012). This could affect the PIC-related gain control system, which is sensitive to serotonergic and noradrenergic inputs, potentially enhancing the net excitatory drive to motoneurons (Lee & Heckman, 1999, 2000). It is also plausible to hypothesise that increased neuromodulatory inputs from brainstem projections (raphe nuclei) primarily converge onto the motoneuron pool on the trained side following unilateral training (Škarabot et al., 2021), potentially explaining the absence of gain in the estimated PIC on the untrained side.

Another possible mechanism underlying the increased ΔF in trained limbs involves changes in the density, number, composition, or function of voltage-gated calcium channels on the motoneuron (Gardiner, 2006). These channels, particularly sodium and L-type calcium channels, are critical for generating PICs that regulate repetitive firing (Binder et al., 2020; Heckman et al., 2005). Evidence suggests that resistance training induces plasticity in the biophysical properties of motoneurons, including alterations in voltage-gated channels, which may enhance the facilitation of spike initiation and the regulation of firing patterns during prolonged exercise (Krutki et al., 2017). Recent studies have also indicated that resistance training may increase the expression of neurotransmitter receptor densities on spinal motoneurons, particularly in those innervating trained muscle fibres, reflecting dendritic plasticity (Adkins et al., 2006; Dai et al., 2024). Interestingly, these adaptations appear activity-dependent, raising the question of cross-education effects, where training one limb induces neural adaptations in the untrained contralateral limb. This phenomenon may be mediated by central mechanisms involving plastic changes in the motoneuron pool, although further research is required to elucidate the underlying neural circuits.

## Limitations And Future Directions

Although HDsEMG enabled the identification of a large population of motor units, it was not possible to directly assess the contribution of spinal and supraspinal mechanisms; instead, it was possible to estimate the contribution of net common inputs converging onto spinal motoneurons. Further studies that concurrently measure brain activity and spinal output are warranted to explore the role of supraspinal, spinal, and peripheral neural determinants governing cross-education.

By calculating ΔF, we were able to estimate training-related effects on this parameter accurately. However, we could not assess sex-based differences in the training response. This limitation is due to the restricted number of participants of different sexes, restricting the statistical power for further exploration. Previous findings (Jenz et al., 2023) have demonstrated baseline differences in ΔF between males and females. Future studies with larger, sex-balanced samples are needed to explore these differences further, thereby providing valuable insights into the role of PICs in muscle force increases and enhancing our understanding of sex-specific training adaptations. Finally, by expanding the understanding of these mechanisms to clinical settings, such as hemilateral muscle and spinal injuries, cross-education would contribute to a more comprehensive aid to significantly improve the quality of life in these populations.

## Conclusions

This study investigated the adaptations elicited by 4-week unilateral resistance training by longitudinally tracking motor units in the biceps brachii of both the exercised and contralateral untrained limbs. Both exhibited significant enhancements in muscle strength and force steadiness following the intervention. Increases in muscle strength were associated with a higher relative shared synaptic input and lower motor unit recruitment thresholds in both limbs. However, larger PIC amplitude and higher motor unit discharge rate were evident only on the exercised side. Improved force steadiness corresponded with a reduced common input variance during steady force production, as well as diminished variability in motor unit spiking activity in both limbs. These findings underscore the contributions of both supraspinal and spinal mechanisms in mediating the transfer of strength and skill from the exercised muscles to the contralateral side. The proportional modifications in neuromechanical variables between the two limbs further support the notion that adaptations in the shared synaptic input implicate motor unit responses facilitating cross-education, with PIC-related adaptations being related to direct mechanical overloading.

## Additional information

### Author contributions

EL and IB conceived and designed the study. EL, PA, and IB collected data. Data analysis and interpretation were collaboratively carried out by EL, PA, ADV, AC, FF, DF, and IB. All authors contributed to manuscript drafting, critical revisions, edits, and the overall refinement of the text. All authors listed meet the criteria for authorship, have made substantial contributions to the work, and have reviewed and approved the final manuscript. All authors agree to be accountable for all aspects of the work.

### Data availability

The data supporting the findings of this study are available upon reasonable request from the corresponding author.

